# Deconvolving mutation effects on protein stability and function with disentangled protein language models

**DOI:** 10.64898/2026.02.03.703560

**Authors:** Kerr Ding, Ziang Li, Tony Tu, Jiaqi Luo, Yunan Luo

## Abstract

Understanding how evolutionary constraints shape protein sequences is fundamental to deciphering the molecular mechanisms underlying protein stability and function, which has broad implications in protein engineering and therapeutics development. Recent advances in protein language models (pLMs) have enabled accurate prediction of mutation effects through evolutionary information, effectively capturing the selective pressure that governs protein sequence variation. A critical challenge, however, remains in disentangling the intertwined mutation effects on protein stability and function, as evolutionary signals conflate both stability-driven and function-driven pressures, obscuring the mechanistic basis of mutation effects and limiting their utility for rational protein engineering. In this work, we introduce DETANGO, a novel deep learning framework that explicitly deconvolves the mutation effects on protein functions by removing components attributable to stability perturbations from the pLM-predicted mutation effects. Guided by computational or experimental stability measurements, DETANGO estimates a functional plausibility score for each single-point mutation that is the component of the mutation effect not accounted for by changes in stability. Single-point mutations with low functional plausibility are predicted to be stable-but-inactive (SBI) variants, whose compromised activities are caused by direct perturbations on functional mechanisms rather than structural stability. Residues enriched for such variants are inferred to be functionally critical, as indicated by the strong evolutionary pressures to maintain protein function. Through extensive benchmarking experiments, we show that DETANGO accurately identifies SBI variants and pinpoints functionally important residues across contexts, including ligand binding, catalysis, and allostery. Moreover, extending DETANGO from individual proteins to homologous protein families reveals shared and distinctive functional patterns across protein families. Collectively, these results establish DETANGO as a biologically grounded framework for disentangling evolutionary constraints on protein stability and function, advancing mechanistic understanding of protein function, and informing rational protein engineering.

## Introduction

Proteins evolve under coupled constraints: their native ensembles must be sufficiently stable to maintain a folded population, while retaining the conformational plasticity required for catalysis, molecular interaction, allosteric signaling, and other functions. Energy-landscape theory formalizes this balance through the principle of minimal frustration, whereby most native residue contacts are cooperatively optimized to favor folding, while permitting localized energetic conflicts where function demands alternative conformations or transiently unfavorable interactions ^1–4^. Mutation effects on stability and function can be concordant, where destabilization reduces abundance and thereby disrupts activity, or be tradeoffs, where activity-enhancing or specificity-shifting substitutions incur modest stability penalties, particularly in locally frustrated neighborhoods near active or binding sites ^5–8^. Recent multidimensional mutagenesis studies ^9–12^ that jointly quantify function and stability readouts of protein variants reveal that, although many mutation effects on function can be explained by stability perturbations, a substantial subset of mutations, especially those located at functional sites, exhibit effects on function far greater than can be accounted for by changes in stability (Fig. 1a).

**Fig. 1.**
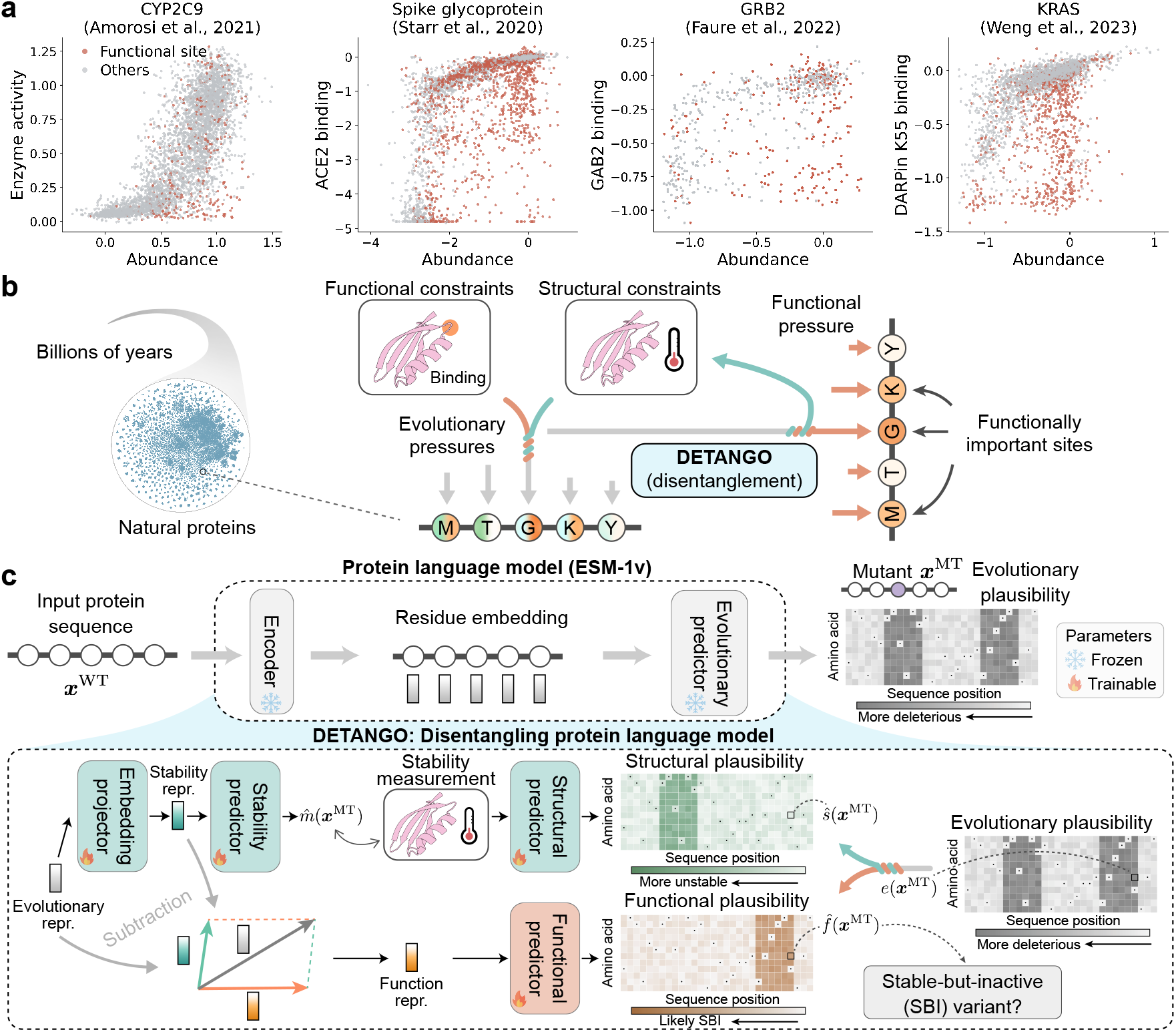
Schematic overview of DETANGO. **a**, Examples of multidimensional mutagenesis experiments ^9–12^ that simultaneously quantify both function readout and stability (cellular abundance) readouts for point-mutation variants. Mutations occurring at function sites annotated in the Conserved Domains Database (CDD) ^47^ are highlighted in red. **b**, Protein residues evolve under intertwined structural and functional constraints. Disentangling function-specific selective pressures from evolutionary pressures is essential for identifying residues critical to protein function. **c**, DETANGO is a deep learning framework that reprograms a pre-trained protein language model (pLM) ESM-1v for deconvolving the mutation effects on protein stability and function. Guided by stability measurements, DETANGO predicts a functional plausibility score for each single-point mutation, reflecting the component of mutation effects attributable specifically to function. repr.= representation. ***x***^WT^ and ***x***^MT^ denote to the wild-type and mutant sequences, respectively. *e*(***x***^MT^), ŝ(***x***^MT^), 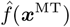, and 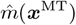 denote the evolutionary plausibility, predicted structural plausibility, predicted functional plausibility, and predicted stability measurement of the mutant, respectively.

Disentangling the intertwined effects of amino acid substitution on stability and function is central to understanding the molecular mechanisms of protein functions. Residues at which most substitutions affect function but not stability, termed ‘stable but inactive’ (SBI) variants ^13^, are often critical for mediating molecular interactions and other functional processes ^11;14;15^. Identifying such functionally important sites is fundamental for decoding the protein sequence-structure-function relationship. While functional residues are typically conserved during evolution, proteins are also subjected to evolutionary constraints to maintain structural stability ^16;17^. Consequently, functional roles inferred from conservation signals alone ^18–22^ may be convoluted with signals that primarily preserve structural stability ^23;24^. Distinguishing sites that directly encode function from those conserved mainly for stability requires a principled disentanglement of mutational effects ^11–13;25^. Likewise, while mutagenesis studies can identify functional sites by probing the effects of amino acid changes on protein function, they often only signal whether a mutation is deleterious, rarely clarifying whether functional loss arises from destabilization of the fold or from direct perturbation of function. Deconvolving these mechanisms is therefore essential for interpreting variant pathogenicity in human genetic diseases ^26^.

Such disentanglement is equally critical for protein engineering. Because beneficial mutations that improve protein function are often destabilizing, engineering campaigns often proceed by coordinating function-enhancing substitutions with compensatory or pre-stabilizing changes ^6;27^. Identifying positions where stability can be effectively traded for function—frequently at locally frustrated regions ^7;28^—again requires resolving the convoluted mutation effects on stability and function. These attributions inform various engineering strategies, including stabilize-then-diversify, targeted exploration of functionally informative yet mutationally tolerant regions, and rational pairing of activity-enhancing and stability-preserving substitutions ^27;29^.

Despite its biological significance, mechanistically disentangling the interplay between stability and function remains a fundamental challenge. Experimental techniques like multiplexed assays of variant effects (MAVEs) quantify changes in molecular phenotypes (e.g., binding, catalytic activity, or cellular growth) induced by mutations ^30;31^. Yet any single phenotypic readout conflates mechanisms: an apparent loss of activity can reflect reduced stability, from altered activity at near-constant stability, or from both ^26^. Multidimensional mutagenesis that jointly measures a function readout with an abundance or stability proxy can partially resolve this ambiguity ^11;12^, but such assays are resource-intensive, system-specific, and remain limited in protein and context coverage. Consequently, a scalable approach that disentangles stability-mediated from function-specific effects is still lacking.

Computational approaches offer a more rapid and scalable alternative to high-throughput mutagenesis studies but face analogous confounding. Conventional analyses combine sequence conservation with biophysical stability estimators and apply heuristic thresholds to separate residues with direct functional roles from positions conserved primarily to maintain the fold ^15;17;25;32–35^. While informative, these post hoc combinations neither intrinsically decompose mutational effects into stability and function nor are they robust to threshold choice. Protein language models (pLMs) have emerged as effective machine learning (ML) tools for predicting mutation effects ^36^ and guiding efficient protein engineering ^37^. Trained to recover partially masked-out residues using the remaining unmasked context, pLMs learn conditional distributions of sequence likelihood from the vast repertoire of protein sequences, thereby capturing the patterns that are likely to occur in natural proteins. The changes in log-likelihood between mutant and wild type have been observed to correlate well with diverse MAVE readouts ^38–42^. However, pLM likelihood changes reflect overall evolutionary plausibility, which aggregates multiple selective pressures; a low-plausibility substitution may be disfavored because it destabilizes the fold, perturbs function, or both ^25;35^. Eliciting functional information from pLMs, therefore, requires an explicit disentanglement of stability-driven and function-specific components.

In this work, we present DETANGO, a reprogrammed pLM that disentangles mutation effects on protein function from those attributable to structural stability. DETANGO leverages the representational capacity of pretrained pLMs and applies a disentanglement operator that removes components explainable by stability perturbations. For a given single-point variant, whereas a standard pLM yields an *evolutionary plausibility* score—changes in sequence likelihood ambiguously reflecting both stability or function—DETANGO estimates a *functional plausibility* score that is the component of the effect on function not accounted for by changes in stability. Variants with low functional plausibilities are predicted as SBI variants, and residues enriched for such variants are inferred to be functionally important.

In benchmark evaluations, DETANGO accurately identifies SBI variants and outperforms post hoc heuristics that combine conservation with stability predictions. The resulting functional-site maps generalize across contexts, including ligand binding (DNA, RNA, peptide, and small molecules), catalysis, and allostery. DETANGO substantially outperforms existing methods that do not disentangle stability from function in functional site identification and, despite being trained without residue-level functional annotations, achieves performance comparable to or even exceeding supervised baselines trained on annotated functional sites ^13;35^. Extending from individual proteins to homologous families, DETANGO reveals both shared and distinctive functional site patterns across protein families, highlighting differentially conserved residues that modulate functional specificity within each family and offering mechanistic insights into functional adaptation of protein families during evolution.

## Results

### DETANGO: disentangling mutation effects on stability and function

DETANGO is a disentangled pLM that explicitly deconvolves the selective pressures shaping protein residues into function- and stability-driven constraints (Fig. 1b). In this work, ‘function’ broadly refers to diverse biochemical activities such as molecular binding, catalysis, and allosteric regulation. DETANGO builds upon recent advances in pre-trained pLMs ^36;37;39;43–46^, whose evolutionary plausibility predictions for protein mutants have been shown to correlate with proteins’ phenotypic readouts from MAVEs and the pathogenicity of human genetic variants. However, when a pLM predicts that a mutation will cause loss of function, it may arise from either reduced structural stability or direct perturbation of functional mechanisms.

To this end, we formalize the disentanglement of mutation effects by positing a multiplicative factorization of the evolutionary probability of a protein sequence ***x*** = (*x*_1_, …, *x*_*L*_) ∈ Σ^*L*^, where Σ denotes the set of amino acids (AAs) and *L* denotes the sequence length, into structural and functional components:

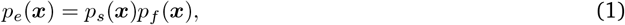

where *p*_*e*_(***x***) denotes the evolutionary probability captured by pLMs as pseudo-likelihood, *p*_*s*_(***x***) the structural probability of ***x***, and *p*_*f*_ (***x***) the functional probability not explained by structural constraints. For each single-mutation mutant sequence ***x***^MT^ of the target wild-type sequence ***x***^WT^, we define its *functional plausibility*, the component of the mutation effect specifically attributable to protein function, as the log functional probability difference between the wild-type protein sequence ***x***^WT^ and the mutant sequence ***x***^MT^:

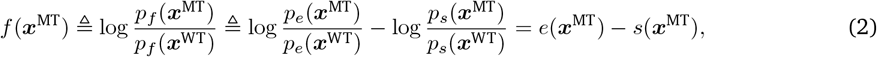

where evolutionary plausibility *e*(***x***^MT^) = log *p*_*e*_(***x***^MT^) − log *p*_*e*_(***x***^WT^) refers to the overall mutation effect predicted by a pLM (Methods) and structural plausibility *s*(***x***^MT^) = log *p*_*s*_(***x***^MT^) − log *p*_*s*_(***x***^WT^) denotes the component attributable to changes in structural stability. Single-point mutations with low functional plausibility scores tend to directly perturb functional mechanisms and thus correspond to SBI variants. Accordingly, residues enriched with such variants are likely to be functionally important residues.

To disentangle the intertwined function- and stability-specific components, DETANGO reprogrammed the pre-trained Transformer-based pLM, instantiated in this work with ESM-1v ^39^, but readily extensible to other state-of-the-art pLMs (Fig. 1c; Methods). Given the wild-type sequence ***x***^WT^ of the target protein as input, DETANGO first decomposes the internal representation of ESM-1v, which captures the evolutionary patterns of proteins, into two latent components: a stability representation and a function representation. The stability representation is explicitly constrained to be predictive of stability measurements (computationally inferred ΔΔ*G* or experimentally assayed cellular abundance), whereas the function representation is obtained by subtracting the stability component from the evolutionary representation. Functional plausibilities are then predicted from these function-specific representations, whereas structural plausibilities are modeled directly from the stability measurements. Unless otherwise specified, DETANGO uses FoldX ΔΔ*G* predictions as the default stability measurements. The training objective of DETANGO enforces that the combination of the structural and functional plausibilities reconstructs the original pLM-predicted likelihood for each single-point mutant, ensuring that the disentanglement remains faithful to the ESM-1v-captured evolutionary signal.

### Accurate classification of stable-but-inactive variants

Mutations that impair protein functions can act through various mechanisms. Some mutations destabilize the protein fold and thereby indirectly compromise activity, whereas other SBI mutations leave stability intact but directly perturb functional mechanisms. Existing mutation effect predictors ^48–52^ do not separate these two mechanisms, labeling all deleterious variants equivalently, despite identifying SBI variants being crucial for elucidating how specific residues contribute to molecular function beyond structural stability. We thus apply DETANGO to systematically identify SBI variants across proteins as a direct evaluation of its disentangling capability.

We assembled a benchmark of MAVEs for 11 proteins from Cagiada et al. ^13^ and ProteinGym v1.0^42^ (Methods), each providing both a functional readout and a stability proxy (cellular abundance) for nearly all single amino acid substitutions. Following Cagiada et al. ^13^, SBI variants were defined as those exhibiting a high cellular abundance (structurally stable) but low functional activity (loss of function; Methods).

As an illustrative example, we applied DETANGO to NUDT15, a human protein that negatively regulates thiopurine activation, where loss-of-function variants increase DNA damage and cytotoxicity. We used the permutation functional plausibility score predicted by DETANGO as the SBI classification score. We compared ground-truth SBI annotations (Fig. 2a) with ESM-1v-predicted mutation likelihood (Fig. 2b), FoldX-predicted thermostability changes (ΔΔ*G*) (Fig. 2c), and DETANGO-predicted functional plausibility scores (Fig. 2d) for all possible point mutations in NUDT15. While ESM-1v broadly classifies most substitutions as deleterious, some of these correspond to highly destabilizing variants (e.g., at residues V17-V19 and C58-A59) with large positive FoldX ΔΔ*G* values, indicating stability-driven loss of function. In contrast, DETANGO disentangles these effects, highlighting substitutions at residues E65, E66, and E112 that preserve structural stability but strongly reduce functional plausibility (Fig. 2d), precisely corresponding to experimentally defined SBI variants (Fig. 2a). Overall, DETANGO’s prediction map more consistently recapitulates the ground-truth SBI landscape than ESM-1v and FoldX.

**Fig. 2.**
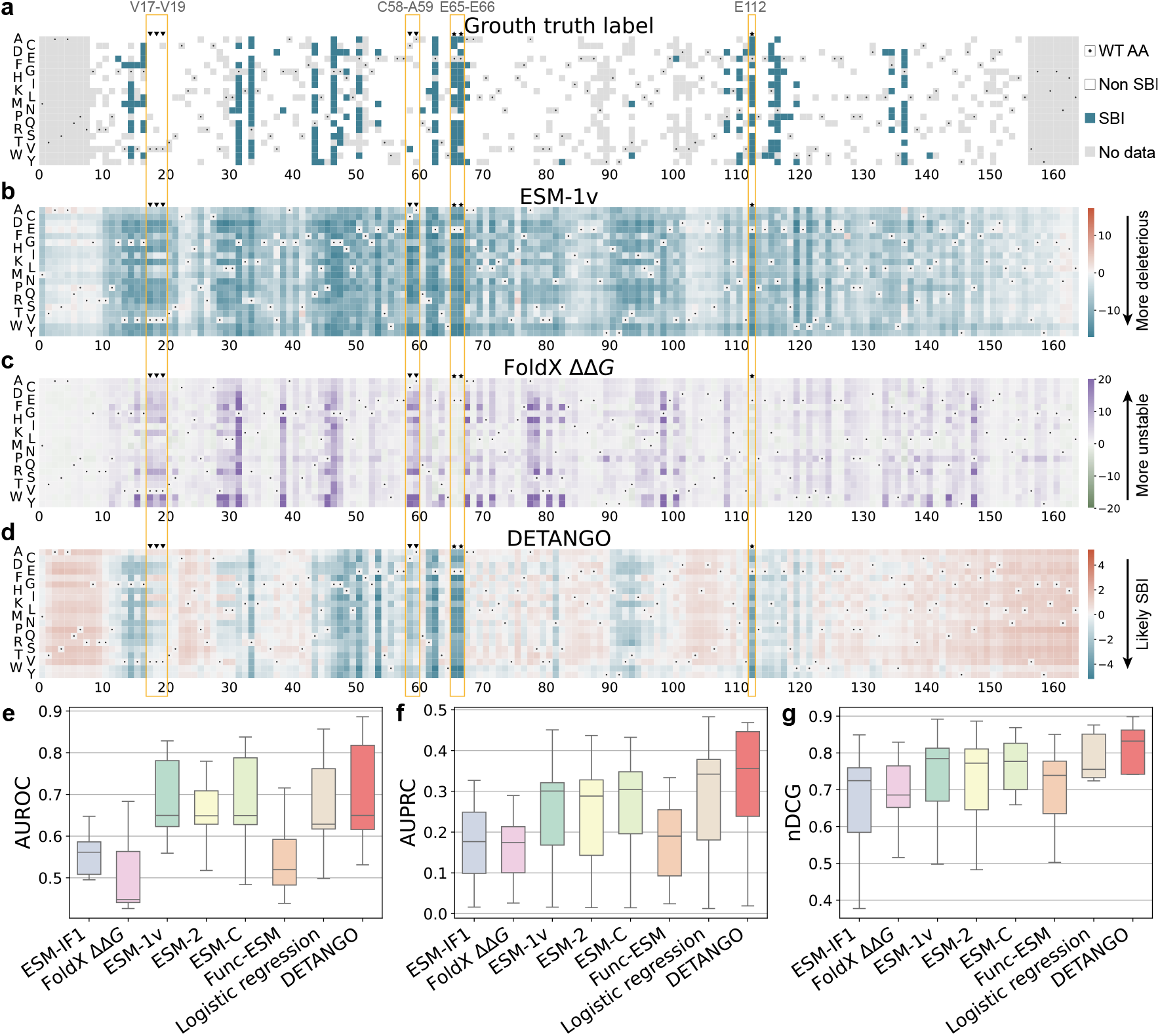
DETANGO accurately classifies stable-but-inactive (SBI) variants in proteins. **a-d**, Heatmaps showing the ground-truth SBI annotations (**a**) and predicted log likelihood from ESM-1v (**b**), FoldX ΔΔ*G* (**c**), and DETANGO functional plausibility scores (**d**) for all single-site mutations in protein NUDT15 (UniProt ID: Q9NV35). Triangles (▼) mark residues enriched in non-SBI variants, while stars (⋆) denote residues enriched in ground-truth SBI variants. **e-g**, Benchmarking results of DETANGO against seven unsupervised baseline approaches (ESM-IF1, FoldX ΔΔ*G*, ESM-1v, ESM-2, ESM-C, Func-ESM, and Logistic regression model) across 11 proteins for SBI variant classification, evaluated by AUROC, AUPRC (**f**), and nDCG (**g**). In box plots, the central line represents the median, boxes indicate the interquartile range (IQR), and whiskers extend to the most extreme data points with 1.5*×*IQR.

This effective disentanglement generalized across the entire benchmark. Across all 11 proteins, DETANGO accurately distinguished SBI from non-SBI variants, achieving consistently better performance across AUROC, AUPRC, nDCG, and F1 metrics (Figs. 2e-g and Supplementary Fig. 1). Existing methods relying solely on stability (FoldX ^53^ and ESM-IF1^54^) or conservation (ESM-1v ^39^, ESM-2^36^, and ESM-C ^55^) performed worse, underscoring the necessity of disentangling function-specific and stability-specific contributions for identifying SBI variants. In addition, DETANGO outperformed heuristic post-hoc approaches such as Func-ESM ^35^ and Logistic regression models ^25;56^ (Supplementary Information A.1), which impose global, context-independent decision boundaries (axis-parallel for Func-ESM and Sigmoidal for Logistic model) across all mutations, positions, and proteins. In contrast, by leveraging contextualized representations from pLMs, DETANGO adapts to local sequence contexts, enabling more accurate SBI detection. Remarkably, although requiring no functional annotations as supervision, DETANGO achieved performance comparable to or even better than that of a supervised method ^13^, which was given advantages by being explicitly trained on three of the 11 proteins in our benchmark (Supplementary Fig. 2).

The strong performance of DETANGO in SBI variant identification highlights its effectiveness to mechanistically disentangle mutation effects. While most existing mutation effect predictions ^48–52^ only provide a binary distinction between benign and deleterious variants, DETANGO offers a refined approach that further separates deleterious variants into those disrupting stability, those perturbing function while remaining stable, and those affecting both, revealing the molecular mechanisms underlying loss-of-function phenotypes.

### Accurate predictions of functionally important sites in proteins

Having validated that DETANGO’s effective disentanglement at the variant level for identifying SBI variants, we next examined whether this capability translates to the residue level, specifically evaluating whether DETANGO can identify amino acid residues directly responsible for protein function, hereafter referred to as functional sites. As previous work has shown that residues harboring more SBI variants are more likely to be functionally critical ^13^, we defined a per-residue *function score* as the negative mean of the functional plausibilities predicted by DETANGO over all possible single substitutions at a given site (Methods). Residues with higher function scores are therefore inferred to contribute more directly to protein function.

We evaluated DETANGO on 408 full-length human proteins from the Human Domainome 1 dataset ^25^ (Methods), which encompasses structurally diverse human domains annotated in the Conserved Domain Database (CDD) ^47^. Among the 213,332 residues in Domainome, 17,381 were labeled as functional sites by CDD, with the remainder considered non-functional. Across all proteins, DETANGO assigned significantly higher function scores to CDD-annotated functional sites compared to non-functional ones (Fig. 3a; *P<*0.001, one-tailed *t*-test). To assess residue-level prediction accuracy within individual proteins, we focused on the AUPRC metric due to the highly imbalanced distribution of functional versus non-functional sites, and also reported AUROC and nDCG. For comparison, we applied several sequence-based pLMs (ESM-{1v, 2, C}) and structure-based models (FoldX and ESM-IF1) to score residues with the same residue-level scoring procedure.

**Fig. 3.**
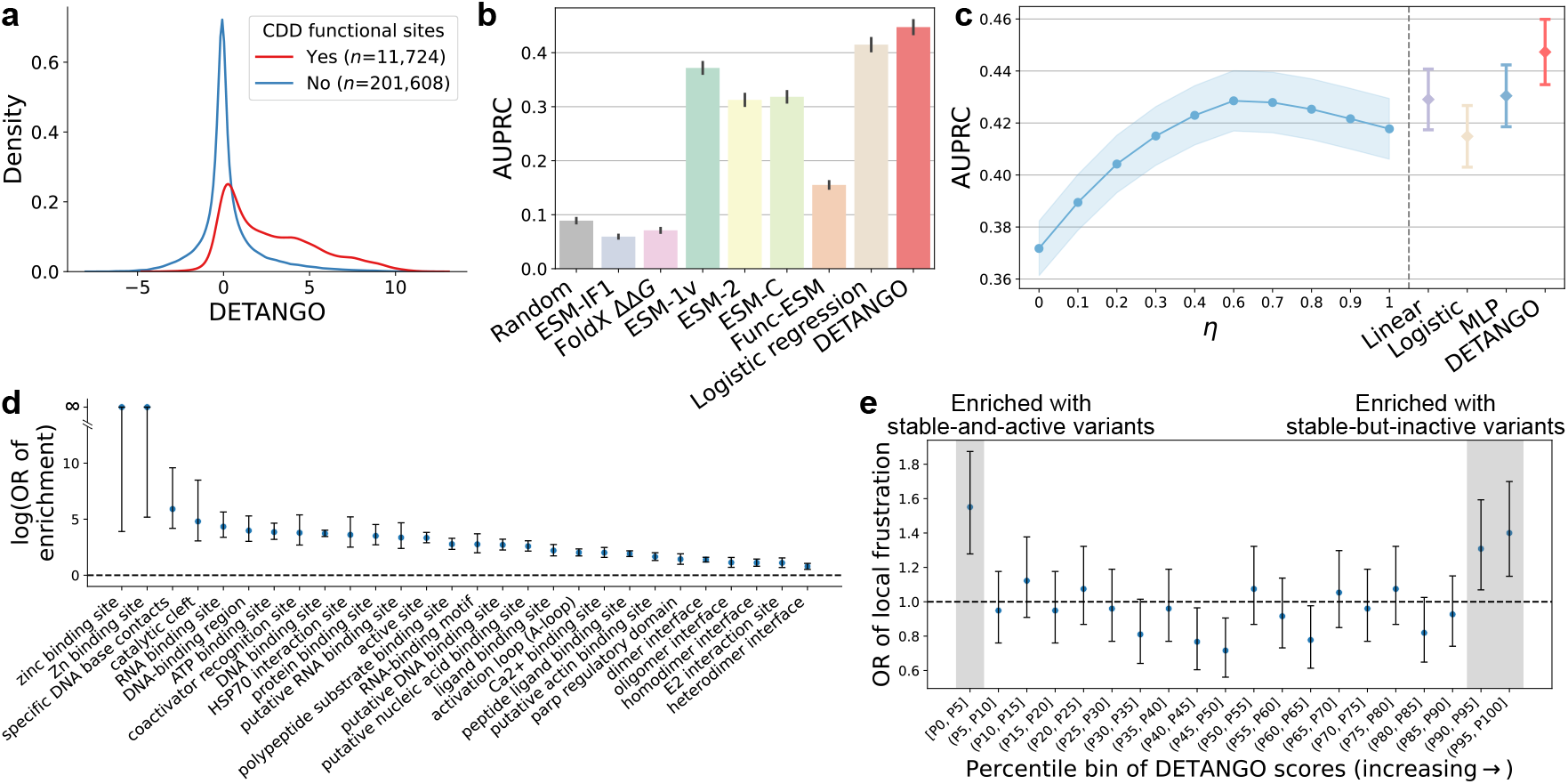
DETANGO accurately identifies functionally important residues across diverse human proteins. **a**, Distribution of residue-level function scores predicted by DETANGO for proteins in Human Domainome 1, comparing residues annotated as functional in the Conserved Domains Database (CDD) with unannotated residues (*P <* 0.001, one-tailed *t*-test). **b**, Performance comparison of DETANGO with seven unsupervised baselines (ESM-IF1, FoldX ΔΔ*G*, ESM-1v, ESM-2, ESM-C, Func-ESM, and Logistic regression model) for identifying CDD-annotated functional residues, evaluated using AUPRC. The AUPRC of a random classifier is shown for reference. **c**, Comparison of DETANGO with four regression-based approaches, including ESM-1v+*η*ΔΔ*G*, Linear regression, Logistic regression, and multi-layer perceptron (MLP) regression, for functional residue identification, evaluated using AUPRC. **d**, Enrichment of major CDD-annotated functional site categories (each containing over 100 annotated residues) among residues with positive function scores assigned by DETANGO. **e**, Enrichment of residues harboring at least two stabilizing mutations across percentile bins of residues ranked by DETANGO from lowest to highest. Residues with high DETANGO function scores are predicted to be functionally important in proteins. Error bars in panels **b–c** indicate the standard error (SE) of the mean, whereas those in panels **d–e** denote 95% confidence intervals (95% CI).

DETANGO achieves the highest accuracy in identifying CDD-annotated functional residues (Fig. 3b and Supplementary Fig. 3). It substantially outperformed its base model ESM-1v and its successors (ESM-2 and ESM-C), likely because those models capture aggregate evolutionary constraints that conflate both function and stability, whereas DETANGO explicitly separates functional signals from aggregate evolutionary information. Interestingly, structure-based models (ESM-IF1 and FoldX) performed near random (Fig. 3b), suggesting that identifying functionally important sites requires quantifying mutation effects beyond stability perturbation alone. DETANGO also outperformed Func-ESM, which infers functional relevance based on the count of SBI variants per residue. Furthermore, a naive linear combination of ESM-1v likelihood score and FoldX ΔΔ*G* (via weighted averaging) performed worse than DETANGO (Fig. 3c; Supplementary Information A.1). When compared to several other methods that identify functional sites by explicitly modeling how much of the aggregate mutation effects (as predicted by ESM-1v) can be explained by changes in stability (as predicted by FoldX) using linear, logistic, or multi-layer perceptron regression, DETANGO consistently achieved higher AUPRC (Figs. 3b-c; Supplementary Information A.1), underscoring the advantage of its representation-level disentanglement over post hoc curve fitting.

We next analyzed DETANGO’s function site predictions for major CDD function annotations (each with more than 100 annotated residues). Enrichment analysis revealed that residues assigned positive function scores by DETANGO were significantly enriched for true functional sites annotated in CDD (Fig. 3d). In particular, all 168 CDD-annotated zinc-binding sites were correctly identified, each receiving a positive function score. We further examined the local energetic frustration at functionally important residues identified by DETANGO. Stabilizing mutations were defined as those with measured cellular abundance values exceeding the 95th percentile of all mutations in Human Domainome 1. Residues harboring at least two stabilizing mutations, which are indicative of strong local energetic frustration, were highly enriched among the top 10% of residues ranked by DETANGO (Fig. 3e), consistent with the established link between functional importance and local energetic conflicts ^3;28^. Interestingly, residues in the bottom 5% of DETANGO predictions also exhibited enrichment for stabilizing mutations (Fig. 3e) and were disproportionately located on solvent-exposed surfaces (RSASA≥0.25, Supplementary Fig. 4), suggesting their engineering potential for enhancing both protein stability and function. As supporting evidence, we observed that such residues with low DETANGO scores were more likely to host stable and active mutations that lead to better-than-wildtype functional activities, as shown by DMS measurements from ProteinGym v1.0 (Supplementary Fig. 5).

Together, these results establish DETANGO as an effective framework for identifying functionally important residues from sequence alone. Whereas historically protein function sites have been found through analysis of sequence conservation that conflates stability and function, DETANGO disentangles these intertwined pressures, revealing residues that are evolutionarily constrained for their roles in function rather than stability preservation.

### Detecting ligand-binding residues in proteins with DETANGO

Building on DETANGO’s demonstrated ability to identify residues broadly important for protein function, we further assessed its capability to pinpoint ligand-binding sites (LBSs). Many proteins perform their biological functions primarily through interactions with ligands, such as small molecules, metal ions, or peptides, at specific binding sites. Accurate identification of these binding residues is essential for understanding molecular mechanisms and for guiding structure-based drug design ^57^. Because mutations at binding interfaces often exhibit larger effects on function than can be explained by stability changes alone ^11^, LBS identification provides an ideal testbed for assessing whether DETANGO’s disentanglement captures functional constraints beyond structural stability.

We evaluated DETANGO as an unsupervised method for LBSs detection on the BioLiP2^58^ database of protein-ligand binding interactions. We used the non-redundant set of BioLiP2 (Methods), which contains 378,814 annotated ligand-binding residues derived from 24,252 experimentally resolved protein structures in Protein Data Bank (PDB) ^59^, spanning a wide range of ligand types, including DNA, RNA, peptide, small molecules, metal ions, and enzyme catalytic residues curated from the Mechanism and Catalytic Site Atlas (M-CSA) ^60^. We measured how well the per-residue function scores predicted by DETANGO align with BioLiP2 annotations using AUPRC. DETANGO consistently outperformed baseline models that either lack disentanglement (ESM-1v) or focus solely on stability prediction (SPURS; Fig. 4a). For evaluation scalability, here we used SPURS, our previously developed ML-based stability predictor ^56^, which is both more computationally efficient and comparably accurate to FoldX for ΔΔ*G* estimation. Notably, DETANGO requires only single-chain sequence and structure inputs, without explicit ligand binding structure, yet achieves robust generalization across diverse ligand types (Fig. 4a), underscoring its scalability and versatility for unsupervised LBS identification.

**Fig. 4.**
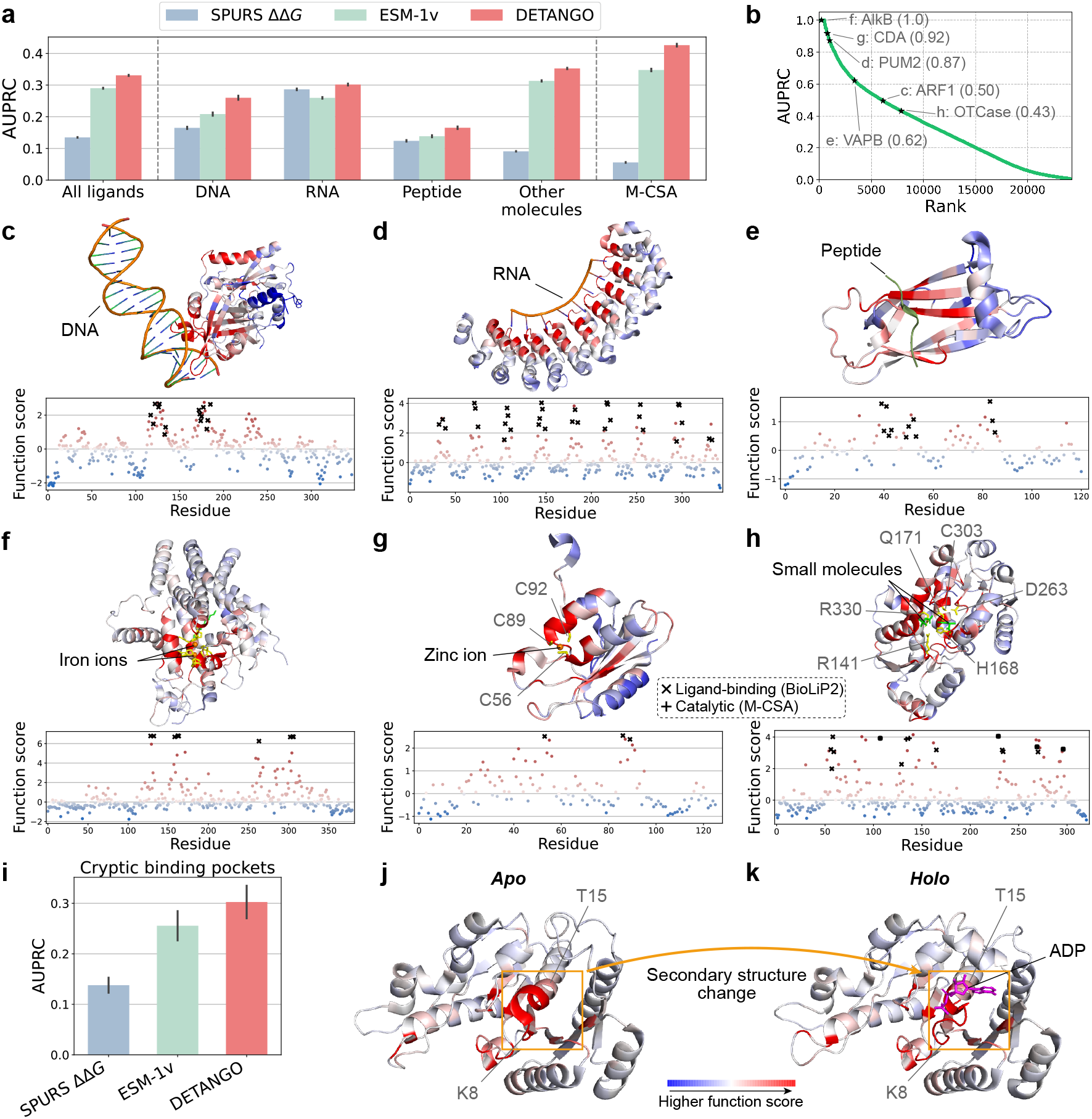
DETANGO identifies ligand-binding and cryptic functional residues in proteins. **a**, Comparison of DETANGO with two unsupervised baselines (SPURS and ESM-1v) for identifying ligand-binding residue in BioLiP2-annotated proteins, evaluated using AUPRC. Average performances across all ligand types (DNA, RNA, peptides, and other molecules) and catalytic residues from the Mechanism and Catalytic Site Atlas (M-CSA) are also shown. **b**, Distribution of DETANGO’s AUPRC performance for identifying ligand-binding residues across 24,252 proteins in BioLiP2. **c-h**, Representative examples of ligand-binding residues identified by DETANGO, including **c**: ARF1 (UniProt ID: Q8L7G0; DNA-binding), **d**: PUM2 (Q8TB72; RNA-binding), **e**: VAPB (Q9QY76; peptide-binding), **f**: AlkB (A0A1I2KHB9; ion binding), **g**: CDA (P9WPH3; zinc binding), and **h**: OTCase (P00480; small-molecule substrate binding). **i**, Comparison of DETANGO, SPURS, and ESM-1v in detecting cryptic binding pockets in proteins from the PocketMiner dataset, evaluated using AUPRC. **j-k**, Example of a cryptic binding pocket identified by DETANGO in CooC1, shown in its *apo* (PDB: 3KJE, chain A) and *holo* (PDB: 3KJG, chain A) states. The identified segment (residues K8–T15) forms an ADP-binding pocket upon structural rearrangement. Residues in structure plots are colored according to DETANGO predictions (red = high functional scores). In scatter plots, BioLiP2 residues are marked with *×* signs (**c**–**h**), and M-CSA catalytic residues with + signs (**h**). Error bars in **a, k** represent standard error (SE) of the mean.

### Case studies of LBS identification

To further illustrate DETANGO’s performance, we examined six representative proteins spanning high to mid AUPRC rankings on the BioLiP2 test set (Fig. 4b). These cases were chosen to represent a range of ligand types (DNAs, RNAs, peptides, small molecules, and metal ions) and structural folds. Mapping DETANGO’s residue-level function score onto 3D protein structures revealed that DETANGO selectively highlights residues enriched for ligand-binding regions.

#### Interactions with macromolecules

For ARF1 from *A. thaliana*, an auxin response factor that binds DNA at auxin response elements ^61^, DETANGO highlighted three clusters of high-scoring residues, two overlapping the DNA-binding domain and one corresponding to its dimer interface (Fig. 4c). A similar pattern was observed for the human Pumilio-2 (PUM2) protein (Fig. 4d), a post-transcriptional repressor ^62^ that binds mRNA through eight alpha-helical repeats arranged in a rainbow-like architecture, each recognizing one base of its RNA target. DETANGO accurately pinpointed the subset of residues mediating RNA interactions across the eight repeats (AUPRC: 0.87). Likewise, in VAPB, an endoplasmic reticulum (ER)-anchored protein involved in ER-mitochondria contact formation ^63^, the top-ranked residues from DETANGO overlapped with the experimentally confirmed peptide-binding residues that interact with the mitochondrial membrane protein MIGA2 (Fig. 4e).

#### Interactions with metal ions and small molecules

In the metalloenzyme alkane monooxygenase (AlkB) from *Fontimonas thermophila*, DETANGO assigned the highest scores to nine histidine residues coordinating two iron ions (Fig. 4f), which have been found critical for the enzymatic functions ^64^. Similarly, in the homotetrameric enzyme cytidine deaminase (CDA) from *M. tuberculosis*, three of the four top-ranked residues by DETANGO (C56, C89, and C92) correspond to zinc-coordinating sites (Fig. 4g). In another small-molecule-binding case of ornithine transcarbamylase (OTCase), an essential liver enzyme for the urea cycle in mammals ^65^, DETANGO precisely localized substrate-binding residues and additionally prioritized residues (R141, H168, Q171, D263, C303, R330) annotated by M-CSA as key catalytic stabilizers of intermediate and transition-state structures (Fig. 4h)

#### Cryptic binding pockets

We further extended our analysis to cryptic binding pockets, i.e., transient ligand-binding sites that are not apparent in static, ligand-free protein structure (“*apo*” form) but emerge upon ligand binding or conformational rearrangement in the ligand-bound state (“*holo*” form) ^66^. These hidden sites are of particular interest because they can render previously “undruggable” proteins amenable to small-molecule targeting, expanding opportunities in structure-based drug discovery. Using the PocketMiner dataset ^67^ (Methods), which catalogs 33 proteins with 760 annotated cryptic residues from PDB, we found that DETANGO’s residue-level function scores were more predictive of cryptic pocket locations than either ESM-1v evolutionary scores or SPURS ΔΔ*G* values (Fig. 4i). A representative example is CooC1 from *C. hydrogenoformans*, a nickel-binding ATPase central to carbon monoxide dehydrogenase maturation ^68^. CooC1 contains a cryptic ADP-binding pocket that is invisible in the *apo*-state structure but becomes evident in the *holo* state during its catalytic cycle. Remarkably, even when provided only with the *apo* structure, DETANGO assigned top function scores to a contiguous segment (residues K8-T15; Fig. 4j) that later forms the ADP-binding pocket through local secondary-structure rearrangement in the *holo* state (Fig. 4k). This example highlights DETANGO ‘s ability to uncover latent functional determinants that are not explicitly encoded in static structures.

Together, these results demonstrated that DETANGO accurately identifies residues mediating diverse ligand bindings using only sequence and single-chain structure as input. By disentangling functional effects from stability constraints, DETANGO generalizes across ligand types and reveals both canonical and cryptic binding determinants, providing a scalable, unsupervised avenue for mapping functional interfaces and guiding structure-based drug design.

### Mapping the allosteric landscapes of proteins

Allostery is the regulation of protein functions through conformational or energetic changes at sites distant from the active sites, representing a fundamental yet incompletely understood mechanism underlying diverse biological processes ^69–71^. Such long-range communication between residues enables coordinated regulation of signaling, catalysis, and macromolecular assembly ^70^. Because allosteric sites often provide higher selectivity than orthosteric (active) sites, they have become increasingly attractive targets for therapeutic intervention ^71;72^. Therefore, the uncovering of allosteric sites holds significant potential for advancing drug design. Having demonstrated that DETANGO can identify orthosteric sites directly involved in ligand binding or other broad functions, we next assessed its capability to map allosteric landscapes across proteins.

We evaluated DETANGO using experimentally characterized allosteric landscapes of KRAS bound to six different binding partners (RAF1, PIK3CG, RALGDS, SOS1, DARPin K27, and DARPin K55) reported by Weng et al. ^12^ (Methods). KRAS is a small GTPase central to cell proliferation and frequently mutated in cancer ^73^. It functions as a molecular switch cycling between inactive GDP-bound and active GTP-bound states through allosteric regulation. Weng et al. ^12^ performed multidimensional deep mutational scanning (DMS) to quantify mutation effects on both protein stability (cellular abundance change; folding ΔΔ*G*_*f*_ ) and function (binding fitness change; binding ΔΔ*G*_*b*_) of KRAS variants. Allosteric sites were defined as distal residues (≥5Å from the binding interface) whose single substitutions yield greater binding perturbation (measured by the average |ΔΔ*G*_*b*_| across possible mutations at that site) than residues at the interface, resulting in 6–22 KRAS allosteric sites (mean=15) per binding partner (Methods). Since this dataset provides experimentally derived cellular abundance measurements that quantify stability changes, we implemented a variant of DETANGO, termed ‘DETANGO (DMS)’, which replaced FoldX-predicted ΔΔ*G* with the experimental stability data to guide disentanglement (Supplementary Information A.2).

We first assessed DETANGO at the mutation level using *in vitro* binding affinity measurements (ΔΔ*G*_*b*_) from Kiel et al. ^74^. Across KRAS variants, per-mutation function scores predicted by DETANGO (DMS) were strongly correlated with experimental ΔΔ*G*_*b*_: variants with more negative DETANGO scores (lower functional plausibility) tended to exhibit higher experimental ΔΔ*G*_*b*_, corresponding to stronger detrimental effects on binding (Fig. 5a). This correlation was markedly stronger than that observed for ESM-1v, confirming the benefit of disentangling function from stability in capturing energetic effects (Fig. 5a). At the residue level, tested on the dataset from Weng et al. ^12^, DETANGO achieved substantially higher AUPRC than ESM-1v and FoldX for identifying allosteric sites across all six KRAS-partner pairs (Fig. 5b). Notably, DETANGO (DMS) outperformed DETANGO (FoldX), highlighting the value of incorporating experimentally measured stability data over computationally predicted scores to inform disentanglement (Fig. 5b).

**Fig. 5.**
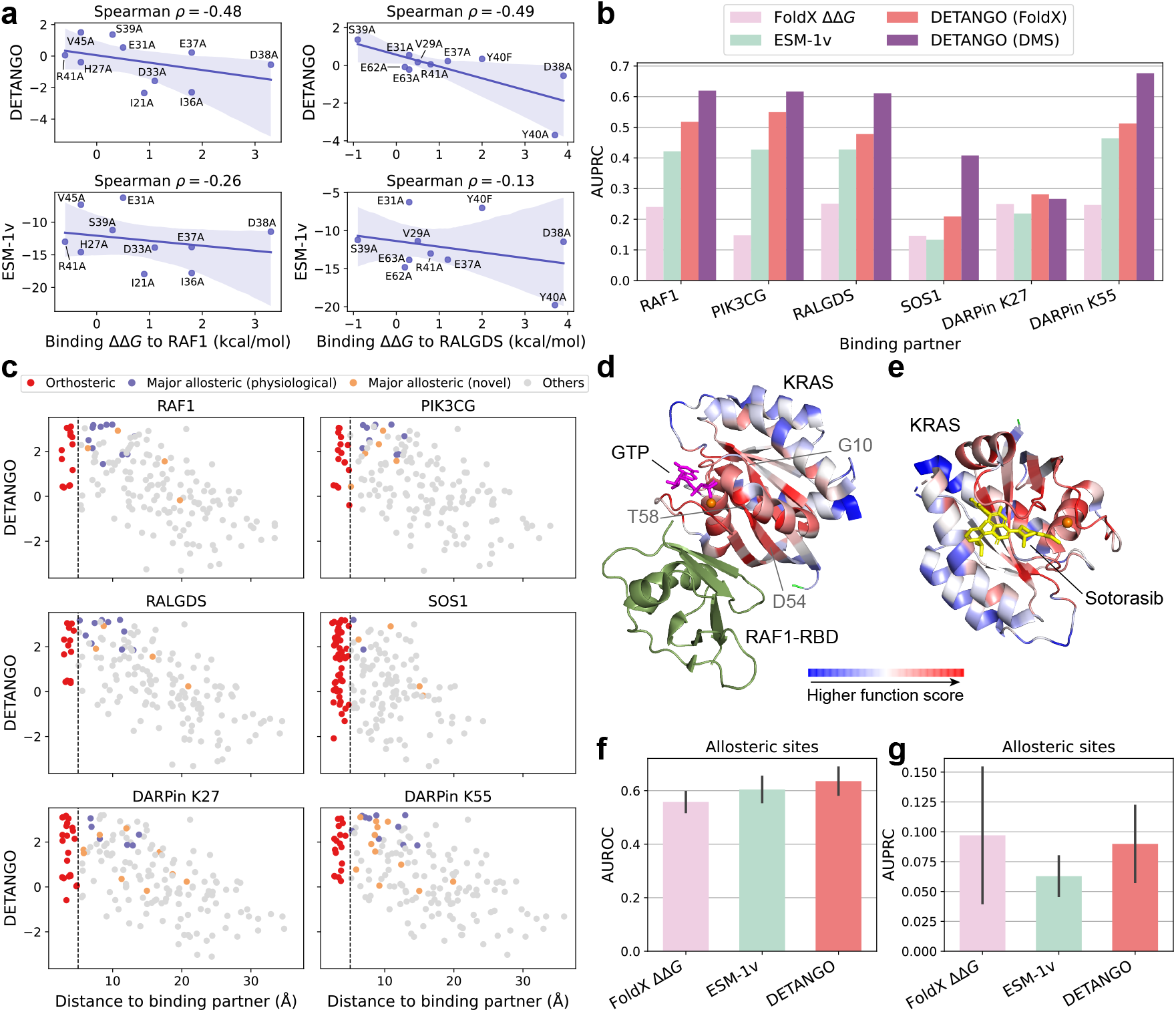
DETANGO reveals allosteric sites in proteins. **a**, Comparisons of DETANGO (DMS) and ESM-1v with *in vitro* binding affinity changes (ΔΔ*G*_*b*_) measured for RAS interactions with RAF1 and RALGDS. Solid lines and shaded areas denote linear regression fits and their 95% confidence intervals, respectively. **b**, Performance comparison of DETANGO (FoldX), DETANGO (DMS), FoldX ΔΔ*G*, and ESM-1v in identifying major allosteric sites of KRAS for each binding partner (RAF1, PIK3CG, RALGDS, SOS1, DARPin K27, and DARPin K55), evaluated using AUPRC. DETANGO (DMS) is a variant of DETANGO that replaces FoldX-predicted ΔΔ*G* with experimentally measured abundance values to guide model disentanglement. The default DETANGO model using FoldX ΔΔ*G* is labeled as DETANGO (FoldX). **c**, Relationship between DETANGO (DMS) function score and residue distance to the corresponding KRAS binding partner (minimal side-chain heavy atom distance). Major allosteric sites are experimentally validated sites whose mutations significantly alter binding free energy and lie at least 5Å away from the interface. Allosteric sites essential for physiological activities are termed physiological, whereas the others are considered novel. **d-e**, 3D structures of KRAS bound to RAF1 (PDB ID: 6VJJ; **d**) and to sotorasib (PDB ID: 6OIM; **e**). KRAS residues are colored based on DETANGO (DMS)’s residue-level function scores, with red indicating higher functional importance. **f-g**, Benchmarking of DETANGO (FoldX), FoldX, and ESM-1v for identifying experimentally validated allosteric sites across 17 proteins, evaluated using AUROC (**f**) and AUPRC (**g**). Bar plots in panels **f-g** represent the mean*±*s.e. of evaluation results over 17 proteins.

Beyond recovering physiological allosteric residues located within conserved nucleotide pockets that bind GTP/GDP, DETANGO identified several novel allosteric sites that significantly influence binding energetics despite lying outside canonical binding interfaces and nucleotide binding pockets (Fig. 5c). For example, three of the four novel allosteric sites (G10, D54, and T58) important for RAF1 binding were assigned with top DETANGO scores (Figs. 5c-d): D54 lies near the binding interface, whereas T58 connects the interface to the nucleotide pocket ^12^ (Fig. 5d). Furthermore, DETANGO successfully prioritized residues forming the binding pocket of sotorasib, a clinically approved allosteric inhibitor targeting KRAS(G12C) for cancer treatment ^75^, suggesting its potential in revealing druggable allosteric regions.

To assess the generalizability of DETANGO beyond KRAS, we extended our analysis to 17 proteins encompassing 159 experimentally validated allosteric sites compiled by Amor et al. ^76^ (Methods). Active-site residue annotations were obtained from BioLiP2, and DETANGO (FoldX) was evaluated for identifying allosteric sites among residues located outside the active sites. DETANGO outperformed FoldX and ESM-1v in AUROC and achieved comparable AUPRC to FoldX, demonstrating robust generalization across diverse protein families. Overall, these results demonstrate the utility of DETANGO for mapping the energetic landscape underlying allosteric regulation and for identifying allosteric sites beyond evolutionarily conserved residues. By decoupling stability-mediated and function-specific effects, DETANGO uncovers how distal residues modulate protein activity through long-range energetic coupling, offering mechanistic insights into allosteric regulation and a scalable approach for discovering previously unrecognized druggable allosteric regions.

### Analyzing conserved functional residues in protein families

Extending beyond individual proteins, we next applied DETANGO to disentangle mutation effects within the evolutionary context of protein families. Motivated by recent evidence that energetic frustration can be conserved across homologous proteins ^28^, we hypothesized that disentangling the intertwined evolutionary pressure, which maintains stability while permitting local unfavorable energetic frustrations for specific functional requirements, could uncover both shared and differential functional patterns among proteins from common ancestry, revealing functional adaptations within divergent protein families.

We applied DETANGO to analyze two representative examples: the mammalian *α* and *β* globin subfamilies and the human RAS superfamily (Methods). In each case, DETANGO derived residue-level function scores for every sequence in the multiple sequence alignment (MSA) of a protein family or subfamily, and scores were averaged across sequences to obtain a function score for each aligned position. Functional sites annotations from the CDD database ^47^ served as ground-truth for evaluation.

In the first case, we analyzed mammalian hemoglobins, including 19 *α* globins and 20 *β* globins, following Freiberger et al. ^28^. Hemoglobins are oxygen-transport proteins in red blood cells, and *α* and *β* globins are parts of the hemoglobin *α*2*β*2 tetramer. CDD annotates 13 and 15 aligned positions in *α* and *β* globins, respectively, as heme binding sites, and 20 positions as the tetramer interface shared by both subfamilies. DETANGO achieved the highest AUPRC for identifying these annotated functional residues compared with FoldX, FuncESM, ESM-1v, and sequence information content (SeqIC), a conservation-based baseline that uses Shannon entropy to quantify MSA variability (Fig. 6a; Supplementary Information A.1). This result demonstrates that disentangling functional signals from evolutionary information can recover functional patterns not captured by raw sequence conservation.

**Fig. 6.**
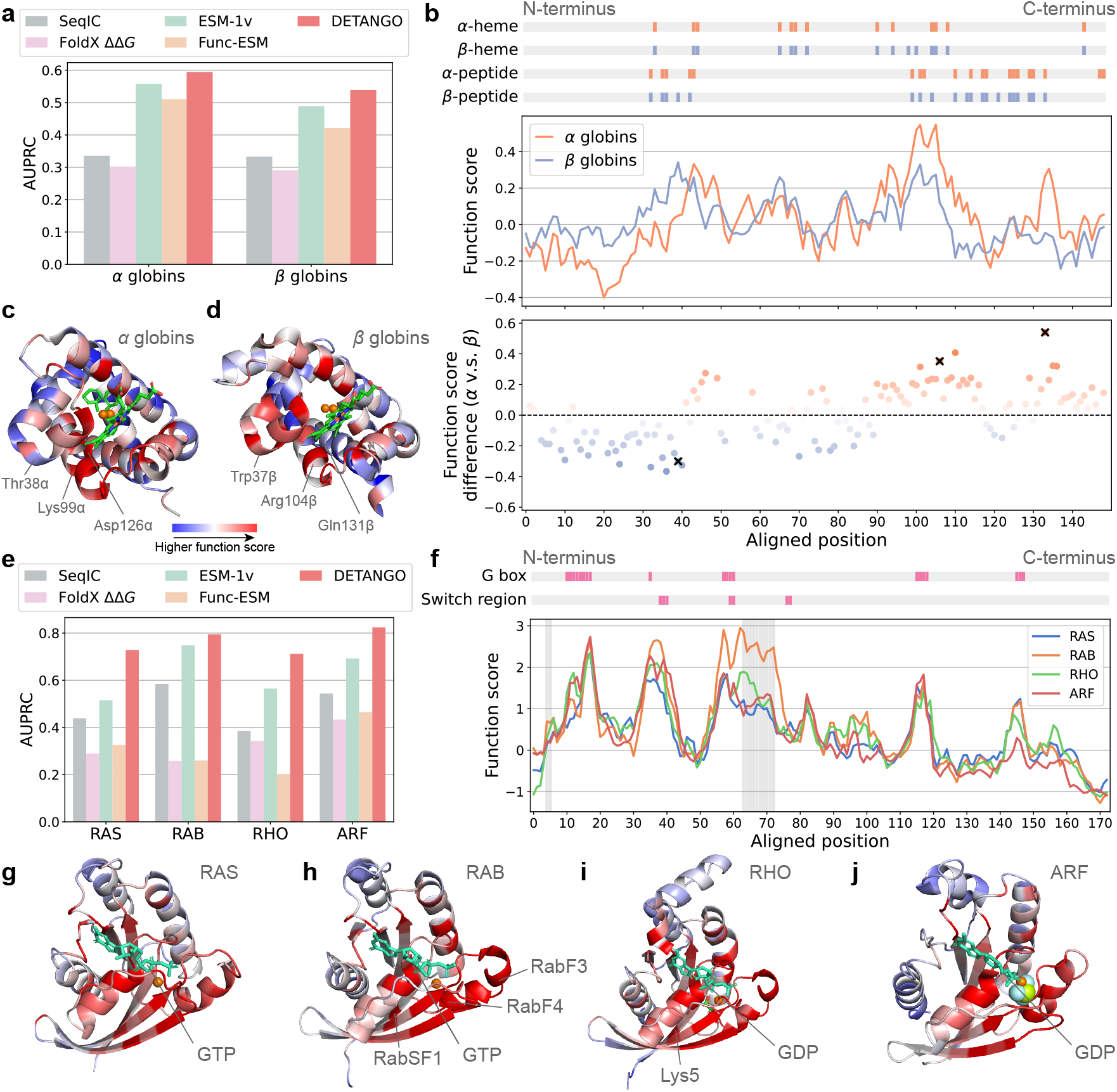
DETANGO reveals shared and differential functional patterns in protein families. **a**, Comparison of DETANGO with four baseline methods (SeqIC, FoldX ΔΔ*G*, ESM-1v, and Func-ESM) for identifying conserved functional sites in *α*- and *β*-globins. **b**, Per-residue function scores predicted by DETANGO for aligned positions in *α*- and *β*-globins. CDD-annotated heme-binding and peptide-binding sites are labeled on top of the plot for reference. **c-d**, Representative structures of *α* (**c**) and *β* (**d**) globins, with residues colored according to DETANGO’s function scores. **e**, Comparison of DETANGO with SeqIC, FoldX ΔΔ*G*, ESM-1v, and Func-ESM for identifying conserved functional sites in RAS, RAB, RHO, and ARF families. **f**, Function scores predicted by DETANGO for aligned positions in the human RAS superfamily. The CDD-annotated G boxes (GTP) and switch regions of the human RAS superfamily are labeled on top of the plot for reference. **g-j**, Representative structures of the RAS (**g**), RAB (**h**), RHO (**i**), and ARF (**j**) families, with residues colored by DETANGO’s function scores. Line plots in panels **b** and **f** were smoothed using a window size of 3.

DETANGO’s per-residue function scores also revealed distinct distribution patterns of functional residues between *α* and *β* globins (Fig. 6b). For example, aligned position 39 (Thr38*α* and Trp37*β*) shows markedly different scores, with Trp37*β* assigned a higher function score than Thr38*α*. This result aligns with CDD annotations labeling Trp37*β* as a tetramer-interface residue whose mutations disrupt intersubunit contacts associated with Trp37*β* at the *α*1*β*2 interface ^77^, whereas Thr38*α* is not annotated as functional by CDD. Conversely, aligned positions 106 and 133, corresponding to Lys99*α* and Asp126*α*, displayed higher scores in *α* globin relative to their *β* globin counterparts. Notably, both positions 106 and 133 have been recognized as specificity-determining positions (SDPs) distinguishing the two subfamilies ^28^. These differences highlight how DETANGO captures the evolutionary fine-tuning of functional constraints following subfamily divergence.

We next applied DETANGO to the human RAS superfamily, a more complex example that includes four major families (RAS, RAB, RHO, and ARF) and plays central roles in signal transduction. Our analyses included 148 human proteins from the RAS superfamily: 37 RAS, 63 RAB, 22 RHO, and 26 ARF family members. CDD annotated 46, 53, 40, and 58 aligned positions in these families, respectively, as family-conserved functional residues. Across all families, DETANGO consistently achieved the best performances in identifying annotated function residues such as GTP binding site and regulator (guanine nucleotide exchange factors (GEF) or guanine nucleotide dissociation inhibitors (GDI)) interaction regions (Fig. 6e). Five prominent peaks in DETANGO ‘s function scores appeared consistently across all four families (Fig. 6f), aligning with the conserved G1–G5 sequence motifs of GTP/GDP-binding sites (Figs. 6f-j). The Switch I/II regions, which undergo conformational changes upon GTP binding, were also assigned with high function scores by DETANGO across all four families (Fig. 6f).

In addition to these shared features, DETANGO uncovered family-specific distribution patterns of functional sites across families. For instance, RAB proteins exhibited significantly elevated function scores at aligned positions 63-72, coinciding with two RAB family motifs, RabF3 and RabF4 ^78^, and at aligned positions 4-5, corresponding to RAB subfamily motif RabSF1^78^ (Figs. 6f, h). These motifs not only serve as hallmarks for RAB family classification but also mediate interaction with Rab escort proteins (REP) ^79;80^, GDP dissociation inhibitors (RabGDI) ^79;80^, and effectors ^81^. Similarly, RHO family proteins were assigned significantly higher function scores at aligned position 5 compared to their counterparts in RAS and ARF families (Figs. 6f, i). These observations are consistent with the CDD annotation of Lys5 as a unique GEF-binding site in the RHO family, which regulates the GTP-GDP exchange reaction ^82^, highlighting a distinctive pattern of functional site distribution within the RHO family.

Collectively, these analyses demonstrate that disentangling mutation effects using DETANGO within the evolutionary context can uncover both conserved and divergent functional determinants across protein families, offering a promising solution to dissect evolutionary constraints shaping protein function.

## Discussion

We introduced DETANGO, a disentangled protein language model that deconvolves the intertwined effects of mutation on protein stability and function. By positing a multiplicative factorization of the evolutionary probability of sequences, DETANGO deconvolves the overall mutation effect into additive components attributable to structural stability changes and to direct disruptions of functional mechanisms. This decomposition reveals a substantial class of stable-but-inactive variants, whose loss of function arises from direct perturbations to function rather than from structure destabilization. Leveraging the function-specific components of mutation effects, DETANGO enables the systematic identification of functionally important residues across diverse biological contexts, including ligand-binding and allosteric sites, and uncovers conserved as well as divergent functional patterns across protein families, which illuminate how proteins adapt their functions over evolution. Although a small subset of residues that jointly contribute to both structural stability and function may not be fully captured by our approach, such cases appear to be relatively uncommon as reported by prior work ^13^, supporting the use of functional plausibility scores as an effective proxy for identifying functionally important residues.

More broadly, this work reframes how evolutionary information can be interpreted to probe the functional roles of protein residues. Traditional approaches infer functional importance of residues primarily from sequence conservation within protein families ^20^, implicitly assuming that residues under strong evolutionary pressure are critical for biological activities. More recently, protein language models (pLMs) have emerged as powerful tools for data-driven estimates of evolutionary probability across protein sequence space ^36;38;39^. However, these evolution-based methods inherently conflate selective pressures arising from coupled constraints to preserve both biological function and structural stability. By explicitly separating function-specific and stability-specific pressures from pLM-derived evolutionary signals, DETANGO enables a more mechanistic interpretation of residue importance beyond conservation alone, offering a principled framework for dissecting how evolutionary constraints shape protein sequences. While contemporaneous work has explored disentangling functional pressures by explicitly contrasting evolutionary energy with folding energy ^83^, DETANGO takes a distinct strategy by learning function-specific representations and directly estimating functional plausibility from them, offering a complementary and representation-driven route for deconvolving functional signals from evolutionary information.

We envision DETANGO as a broadly applicable tool for proteome-scale biological discovery. Requiring no residue-level functional annotations during training, DETANGO can be readily applied to the rapidly expanding set of proteins with known sequences and high-confidence structures predicted by AlphaFold ^84^. Computational analyses on BioLiP2 show that DETANGO reliably identifies residues involved in intermolecular interactions with diverse ligands, including DNA, RNA, peptides, metal ions, and small molecules. Accurate detection of such ligand-binding sites is central to structure-based drug discovery, where candidate compounds are virtually screened against functional pockets to guide therapeutic design ^85^. Beyond annotation, functional residues prioritized by DETANGO provide a rationale starting point for protein engineering, enabling the design of sequences that scaffold key functional sites to recapitulate or modify target functions ^86;87^. In addition, whereas existing variant pathogenicity predictors typically produce a single conflated score ^41;52;88^, DETANGO offers the possibility of mechanistic interpretation of pathogenic missense variants by distinguishing effects driven by structural destabilization from those arising from direct disruption of function.

Although demonstrated here using ESM-1v, the applicability of DETANGO is not tied to a specific pLM. As a general framework for disentangling evolutionary information encoded in pLMs, DETANGO can be extended to recent expressive biological foundational models ^36;55;89^. Moreover, the stability component integrated into DETANGO can be replaced with alternative sources of stability information, whether experimental or computational. Similarly, pLM-predicted evolutionary plausibilities can be substituted with functional readouts from deep mutagenesis studies, enabling interpretable disentanglement of mutation effects directly from experimental data. This flexibility underscores the versatility of DETANGO as a general tool for interrogating the mechanistic basis of protein variants.

This work also contributes to the growing efforts to interpret the biological knowledge embedded in pLMs, examining how learned representations align with structural and functional aspects in proteins ^90–98^. Recent advances in mechanistic interpretability have shown that sparse autoencoders (SAE) can decompose pLM representations into interpretable residue-level features that correlate with functional annotations ^91^, regulatory motifs ^89^, structural features ^89;90;92^, and conserved domains ^90;92^, as well as protein-level features corresponding to protein families ^91^. Rather than probing representations directly, DETANGO demonstrates that evolutionary signals captured by pLM predictions can be explicitly decomposed into mechanistically meaningful components. While the present study focuses on disentangling stability- and function-related effects, the framework is inherently extensible and can be generalized to isolate finer-grained biochemical constraints encoded in evolutionary data. Expanding DETANGO along these directions offers a principled path toward a deeper understanding of how pLMs internalize evolutionary constraints and functional features, with important implications for protein science, evolutionary biology, and protein engineering and design.

## Methods

### Datasets and experimental setup

#### Stable-but-inactive (SBI) variants

To evaluate the capability of DETANGO to classify SBI variants, we collected the multidimensional DMS for 11 proteins, each including both a functional readout and a cellular abundance readout for single amino acid substitutions. Cellular abundance measurements have been shown to correlate with protein stability ^99;100^. The MAVEs for four proteins, including NUDT15, PTEN, CYP2C9, and GRB2, were compiled by Cagiada et al. ^13^, while the MAVEs for the remaining seven proteins, including GCK, KCNE1, DAO1, HLA-A, KRAS, SLC22A1, and VKORC1, were retrieved from the ProteinGym v1.0 benchmarking dataset ^42^. Data sources for all included MAVEs are summarized in Supplementary Table 1. Following Cagiada et al. ^15^, raw DMS assay scores were binarized by fitting a three-component Gaussian mixture model and defining the cutoff as the intersection between the first and third Gaussian distributions. Binarized scores of 0 and 1 indicate low and high readouts, respectively. Consistent with Cagiada et al. ^13^, SBI variants were defined as variants with low functional readout (0) and high cellular abundance readout (1).

For stability prediction, FoldX ΔΔ*G* and ESM-IF1 scores were computed using AlphaFold2-predicted structures ^84^. Structures were retrieved from AlphaFold DB ^101^, and for proteins not available in the database, structures were predicted using ColabFold ^102^.

#### Conserved Domains Database (CDD)-annotated functional sites

Human Domainome 1^25^ is a large-scale reference dataset that quantifies the cellular abundance for over 500,000 human protein variants spanning structurally diverse domains across multiple protein families. The dataset covers approximately 2.1% of all human proteins, 1.2% of all domains, and 2.0% of unique domain families in the human proteome. Functional residue annotations for proteins in Human Domainome 1 were retrieved from the Conserved Domains Database (CDD) ^47^ via the InterPro API ^103^. Among 563 proteins in Human Domainome 1 with available CDD annotations, 408 pass ESM-1v’s sequence length constraint (≤1,022 AAs). Across these proteins, 213,332 residues were analyzed, of which 17,381 were annotated as functional sites by CDD, with the remainder considered as non-functional. AlphaFold2-predicted structures ^84^ were used for FoldX ΔΔ*G* calculations, retrieved from AlphaFold DB ^101^ when available or generated using ColabFold ^102^ otherwise.

For analyses of local energetic frustration, stabilizing mutations were defined as variants whose measured abundances exceeded the 95th percentile of all mutation abundances in Human Domainome 1 (threshold = 0.1275). To assess the engineering potential of residues with the lowest DETANGO scores, we analyzed three DMS assays from ProteinGym v1.0 (DLG4_HUMAN_Faure_2021, DLG4_RAT_McLaughlin_2012, and GRB2_HUMAN_Faure_2021) that evaluate variant functions for proteins represented in Human Domainome For each protein, we compared the abundance and functional measurements of variants at residues ranked in the bottom 10% by DETANGO against all other residues (Supplementary Fig. 5).

#### Ligand-binding sites (LBSs)

We evaluated DETANGO for LBS detection using BioLiP2^58^ and for cryptic binding pockets detection using PocketMiner ^67^. BioLiP2 is a semi-manually curated database derived from Protein Data Bank structures ^59^, providing high-confidence annotations of residues in proteins that bind to biologically relevant ligands, including DNA, RNA, peptide, small molecules, and metal ions. In addition, BioLiP2 incorporates enzyme catalytic residues through mapping from the Mechanism and Catalytic Site Atlas (M-CSA) ^60^.

We used the non-redundant dataset of BioLiP2, where ligand-binding proteins share less than 90% sequence identity as calculated by CD-HIT ^101^. After filtering out ligand-binding proteins with sequences longer than 1,022 AAs, containing non-canonical AAs, or exhibiting missing residues in the PDB, the final evaluation set comprised 24,252 proteins with 378,814 annotated ligand-binding residues.

PocketMiner ^67^ contains 38 curated pairs of *apo*-*holo* protein structures filtered from the PDB that cover a diverse space of cryptic pocket structures and conformational changes. There are 39 cryptic binding pockets in total that have large deviations between the *apo* and *holo* states of proteins. The structural motions between the *apo* and *holo* states of protein structures include loop motion, secondary structure motion, secondary structure change, and interdomain motion. After filtering out proteins with sequences longer than 1022 AAs or having missing residues in the PDB, we evaluated DETANGO on 33 proteins from PocketMiner, encompassing 760 annotated cryptic binding residues. For SPURS ΔΔ*G* prediction, single-chain PDB structures were used as input.

#### Allosteric sites

To assess the capability of DETANGO for mapping the allosteric landscapes of proteins, we first used the experimentally derived allosteric landscapes of KRAS binding to six binding partners (RAF1, PIK3CG, RALGDS, SOS1, DARPin K27, and DARPin K55) ^12^ for evaluation. Multidimensional DMS was performed to quantify both the cellular abundance of KRAS mutants and their binding fitness to the six binding partners. MoCHI ^104^ was applied to decompose the raw fitness measurements of single-mutation KRAS variants into folding free energy changes (ΔΔ*G*_f_) and binding free energy changes (ΔΔ*G*_b_).

For each binding partner, active site residues were defined as those within 5Å of the binding partner (minimal side-chain heavy-atom distance). Allosteric sites were defined as distal residues at least 5Å) from the binding partner whose single-point mutations yield a higher average absolute binding ΔΔ*G* than active-site residues. Under this definition, KRAS contains 15, 15, 14, 6, 15, and 22 allosteric sites in its interactions with RAF1, PIK3CG, RALGDS, SOS1, DARPin K27, and DARPin K55, respectively. To further test its generalizability for identifying allosteric sites, we evaluated DETANGO on a curated dataset of 20 allosterically regulated proteins compiled by Amor et al. ^76^. Active-site annotations were obtained from BioLiP2, and allosteric site identification was evaluated among residues outside active sites. After filtering out proteins that do not have active sites annotated in BioLiP2, 17 proteins with allosteric site annotations were retained in the test set.

#### Functional patterns in protein families and superfamilies

We analyzed the mammalian globin family and the human RAS superfamily as examples to demonstrate the effectiveness of DETANGO in uncovering conserved and differential functional patterns among protein families, following Freiberger et al. ^28^. Mammalian hemoglobins are oxygen-transport proteins in red blood cells, consisting of *α* and *β* globin subunits assembled into the hemoglobin *α*2*β*2 tetramer. For *α* and *β* globins of mammals, we used 20 hemoglobins collected by Freiberger et al. ^28^ from the PDB by April 2022. CDD annotates 13 and 15 aligned positions in *α* and *β* globins, respectively, as heme-binding residues, along with 20 sites as the tetramer interface shared by both subfamilies.

Following Freiberger et al. ^28^, we retrieved the human proteins belonging to the RAS superfamily from Rojas et al. ^105^. Excluding proteins with deprecated or unmatched UniProt identifiers ^106^ and the RAN family containing only one protein, the final dataset comprised 148 proteins that belong to four protein families: RAS (n=37), RAB (n=63), RHO (n=22), and ARF (n=26). CDD annotated 46, 53, 40, and 58 aligned positions in these families, respectively, as family-conserved functional residues. Multiple sequence alignments of proteins belonging to the same superfamily or family were generated using MAFFT ^107^ with default parameters. Details of the analyzed proteins in the globin family and human RAS superfamily are provided in Supplementary Tables 2 and 3. AlphaFold2-predicted structures were used for protein stability prediction.

### Computations of evolution-based scores and stability scores

#### ESM-1v for evolution-based mutation effect prediction

ESM ^36;37;45^ is a series of protein language models (pLMs) trained on the UniRef protein sequence database ^108^ using a masked language modeling objective, enabling it to capture evolutionary constraints from millions of natural protein sequences. Let a protein sequence be denoted as ***x*** = (*x*_1_, …, *x*_*L*_) ∈ Σ^*L*^, where Σ is the set of amino acids (AAs) and *L* is the sequence length. Given the sequence context ***x***−*i* = (*x*_1_, …, *x*_*i*−1_, *x*_*i*+1_, …, *x*_*L*_) that masks residue *x*_*i*_, ESM predicts the conditional probability *p*(*x*_*i*_|***x***_−*i*_) of observing amino acid *x*_*i*_ at position *i*. For a protein variant with a single-site mutation at position *t*, ESM can be used to score its mutation effect, referred to as evolutionary plausibility in this work, using the log-odds difference between the mutant sequence ***x***^MT^ and the wild-type sequence ***x***^WT^:

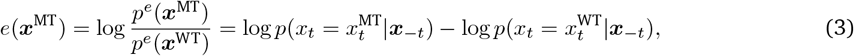

where ***x***_−*t*_ denotes the sequence ***x*** with residue *t* masked. In this work, we instantiated the pLM in DETANGO using ESM-1v, motivated by its strong empirical performance in mutation effect prediction ^39^, but it can be readily extended to more recent ESM models ^36;37^. The final evolutionary plausibility score *e*(***x***^MT^) is computed by averaging predictions from five individual pre-trained ESM-1v checkpoints (i.e., esm1v_t33_-650M_UR90S_1 to esm1v_t33_650M_UR90S_5). Variants with *e*(***x***^MT^) *<* 0 are considered less evolutionarily plausible than the wild type. For residue-level embedding extraction, we used the final-layer embedding of esm1v_t33_650M_UR90S_1 for each residue. This choice preserves generalizability while substantially reducing GPU memory requirements, enabling scalable embedding extraction without the prohibitive GPU memory overhead associated with the loading and inference of all five ESM-1v checkpoints simultaneously.

#### Protein stability prediction

Protein stability change upon sequence mutations was quantified using ΔΔ*G*, defined as the change in Gibbs free energy change between the wild-type (WT) and mutant (MT) structures. FoldX ^53^ was used to compute ΔΔ*G* for protein variants in the SBI dataset, Human Domainome 1, the allosteric datasets, and the family-level analyses. Prior to ΔΔ*G* computation, protein structures were first processed using the FoldX “RepairPDB” command, followed by mutation modeling with the “BuildModel” command.

For large-scale stability prediction for the variants of 24,252 proteins in BioLiP2 and 33 proteins in PocketMiner, we used SPURS ^56^, our recently developed deep learning-based protein stability predictor that offers substantially higher computational efficiency than FoldX while achieving state-of-the-art predictive performance for protein stability ^56^. Given a single-chain protein structure as input, SPURS predicts ΔΔ*G* values for all single-site mutations in a single forward pass, requiring less than one minute on a single NVIDIA A40 GPU. By comparison, FoldX typically requires approximately 20 minutes on 60 CPU cores to compute ΔΔ*G* for all single-site mutations of a single protein.

#### Integrative baseline approaches combining evolutionary and stability scores

In addition to methods that rely exclusively on either evolutionary or stability predictions, we evaluated a set of integrative baseline approaches that jointly incorporate both sources of information. These included Func-ESM ^35^, Logistic regression model ^25;56^, and a supervised Functional model ^13^. We additionally implemented linear regression, MLP regression, and a weighted averaging scheme that combines ESM-1v evolutionary plausibility scores with FoldX ΔΔ*G*. Detailed descriptions of all baseline implementations are provided in Supplementary Information A.1.

### DETANGO: pLM reprogramming to deconvolve mutation effects

#### Prediction of functional plausibility

To model the function-specific component *f* (***x***^MT^) of mutation effects, DETANGO introduces a disentanglement framework that decomposes the internal representation of a pre-trained pLM (ESM-1v) for an input wild-type protein sequence, which captures aggregated evolutionary constraints, into two latent components: a stability representation and a function representation (Fig. 1c). Specifically, given a wild-type protein sequence ***x***^WT^, the encoder of ESM-1v maps residue at position *t* to an evolutionary embedding 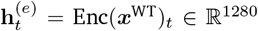, where Enc(·) denotes the encoder. DETANGO decomposes **h**^(*e*)^ into a function-specific representation 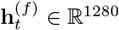 and a stability-specific representation 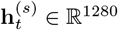 for the *t*-th residue of the wild-type protein sequence, such that

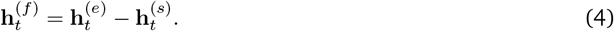

The stability-specific embedding 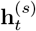 is obtained by projecting 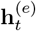 through a multi-layer perceptron (MLP)-based projection network 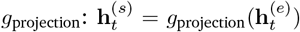. To ensure that 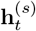 captures information relevant to protein stability, 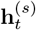 is further passed to an MLP-based stability prediction network *g*_stability_, which predicts stability measurements (e.g., ΔΔ*G* or cellular abundance) for all single-point mutations at position *t*:

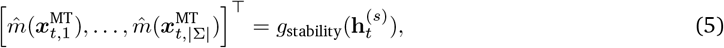

where 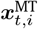 denotes the mutant sequence in which residue *t* is substituted by the *i*-th AA in the alphabet Σ. While the separation between stability- and function-specific representations is operational rather than strictly guaranteed, our formulation aims to isolate components of the evolutionary signal that are preferentially associated with stability versus with functions.

After decomposition, the function-specific embedding 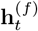 is used to predict the functional plausibility of all possible single-point mutations at position *t* through an MLP-based functional predictor *g*_functional_:

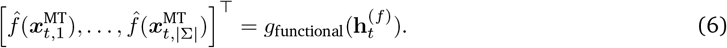

#### Prediction of structural plausibility

DETANGO models the stability-driven component *s*(***x***^MT^) of mutation effects using either computationally predicted ΔΔ*G* values (FoldX ^53^ or SPURS ^56^) or experimentally measured cellular abundance values, denoted as *m*(***x***^MT^). Under a Boltzmann distribution, the stability-specific probability of a protein sequence ***x*** follows 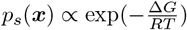, where Δ*G* is Gibbs free energy change, *R* is the gas constant, and *T* is the thermodynamic temperature. Accordingly, the structural plausibility *s*(***x***^MT^) of a mutant sequence relative to the wild type can be expressed as a function of ΔΔ*G*:

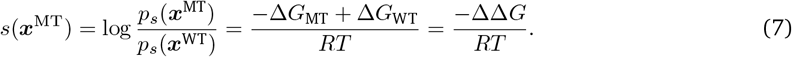

Rather than imposing a fixed functional form, DETANGO uses a lightweight MLP, *g*_structural_, to learn a data driven mapping between the stability-relevant measurement *m*(***x***^MT^) and the corresponding predicted structural plausibility ŝ(***x***^MT^) for a single-point mutant:

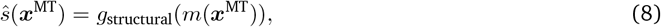

This formulation allows DETANGO to flexibly incorporate stability-relevant metrics beyond ΔΔ*G* and cellular abundance without modifying the core framework.

#### Objective function

To ensure that the sum of predicted structural and functional plausibilities reconstructs the original evolutionary plausibility score produced by the pLM, DETANGO minimize a disentanglement loss defined as the mean squared error (MSE) between the original ESM-1v score and the sum of the decomposed scores for all single-point mutations at each residue *t*:

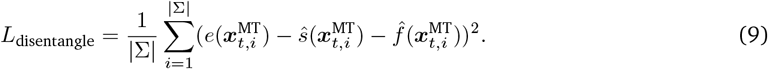

To encourage the stability-specific embedding to be predictive of stability measurements, an additional MSE loss is applied to jointly train *g*_projection_ and *g*_stability_:

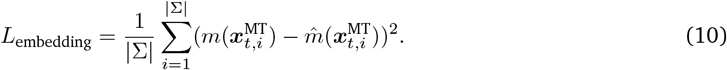

To mitigate model overfitting and prevent trivial solutions, an *L*_2_ norm regularization is applied to the predicted functional plausibilities: 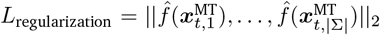. When stability measurements (e.g., abundance) are unavailable for certain mutations, a masking strategy is applied so that loss terms are computed only over mutations with known measurements (Supplementary Information A.2). The full loss function for training DETANGO is given by:

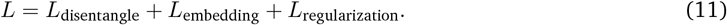

We additionally explored incorporating a mutual information regularization term *L*_MI_ to further promote embedding disentanglement, but observed only marginal improvements and therefore did not include it in our final model (Supplementary Information A.2).

#### Training details

DETANGO requires no explicit residue-level functional annotations as supervision. For each protein in the dataset, we individually trained DETANGO using the loss function described above. Hyperparameters were selected by minimizing validation losses (Supplementary Information A.2). Training was performed using the AdamW optimizer with default parameters, with a learning rate of 5 *×* 10^−4^, a batch size of 16, and up to 100 training epochs, with early stopping applied if validation loss failed to improve for five consecutive epochs.

The hidden dimensions of *g*_structural_, *g*_functional_, and *g*_stability_ were set to 256, each implemented as two-layer MLPs. The projection network *g*_projection_ used a hidden dimension of 64 with three linear layers. All MLP layers were followed by GELU activations, dropout with probability 0.1, and layer normalization. Models were trained on a single NVIDIA A40 GPU. For each protein, we performed a 10-fold cross-validation, where residues are partitioned into training and validation sets across folds, and constructed the final DETANGO model as an ensemble of the 10 independently trained replicas. Training a single model required less than one minute, with training the full ensemble completed in under ten minutes.

#### Identification of functionally important residues

To quantify the functional importance of the *t*-th residue in a protein sequence, DETANGO examined the functional plausibility of all possible single-point mutations at that position. We define the residue-level *function score* of the *t*-th residue as 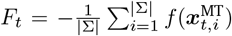. A higher residue-level function score indicates that the residue is more likely to be enriched with mutants that directly perturb function but not stability, indicating a more important role in the protein’s biological function.

## Data availability

This study used publicly available protein fitness and function datasets for computational evaluation: Cagiada et al. ^13^ (https://github.com/KULL-Centre/_2022_functional-sites-cagiada); ProteinGym benchmark v1.0 (https://github.com/OATML-Markslab/ProteinGym); Human Domainome 1 (https://github.com/lehner-lab/domainome); BioLiP2 (https://zhanggroup.org/BioLiP/index.cgi); PocketMiner (https://www.nature.com/articles/s41467-023-36699-3#data-availability); energetic landscape of KRAS (https://github.com/lehner-lab/krasddpcams); Amor et al. ^76^ (https://www.nature.com/articles/ncomms12477); mammalian globin family and the human RAS superfamily (https://doi.org/10.5281/zenodo.10093060); Conserved Domain Database (CDD) annotations (https://www.ncbi.nlm.nih.gov/cdd/). AlphaFold2-predicted protein structures (v4) were obtained from the AlphaFold DB (https://alphafold.ebi.ac.uk/), while experimentally determined structures were retrieved from the Protein Data Bank (https://www.rcsb.org/).

## Code availability

The source code of DETANGO is available at https://github.com/luo-group/DETANGO. DETANGO was built on Python 3.9, PyTorch 2.2.0, NumPy 1.26.4, Pandas 2.2.3, Biopython 1.84, Scikit-learn 1.4.2, SciPy 1.13.1, and Transformers 4.34.0. ESM-1v (https://github.com/facebookresearch/esm) was used for evolutionary plausibility prediction. FoldX5 (https://foldxsuite.crg.eu/) and SPURS (https://github.com/luo-group/SPURS) were used for thermostability prediction. ColabFold (https://github.com/sokrypton/ColabFold) was used to predict protein structures for sequences lacking AlphaFold2-predicted models in the AlphaFoldDB. MAFFT (https://mafft.cbrc.jp/alignment/server/index.html) was used to align protein sequences using default parameters.

## Acknowledgments

This work is supported in part by the National Institute of General Medical Sciences of the National Institutes of Health (R35GM150890), a National Science Foundation (NSF) CAREER Award (No. 2442063), an NSF ACED grant (No. 2435754), and the NSF Molecule Maker Lab Institute funded by the NSF under Awards No. 2019897 and 2505932. The authors also acknowledge the CloudHub GenAI Seed Grant funded by GaTech IDEaS and Microsoft. We thank Haoyu Wang (Georgia Institute of Technology) for insightful discussions on the conceptual development and implementation of this study.

## Author contributions

Y.L. conceived and supervised the research project. K.D. and Y.L. developed the computational method. K.D. implemented the software and performed the computational experiments. K.D., Z.L., and T.T. curated the data for benchmarking experiments. K.D. analyzed the benchmarking results of the computational model with support from Y.L. and Z.L. K.D. and J.L. performed visualization of the benchmarking results with support from Y.L. K.D., Y.L., and J.L. wrote the manuscript with support from all authors.

## A Supplementary Information

### A.1 Implementation of baseline approaches

We assessed the performance of DETANGO for identifying stable-but-inactive (SBI) variants and detecting functionally important residues in proteins. For reference, we included a variety of baseline approaches in benchmarking experiments. Evolution-based predictors and stability-based predictors were included to demonstrate the necessity and effectiveness of disentanglement. We further included integrated approaches that combine evolutionary information and stability information for identifying functionally important protein residues.

#### Evolution-based predictors

Trained on natural protein sequences, protein language models (pLMs) learn to capture the evolutionary patterns of proteins. They can be leveraged to identify evolutionarily implausible variants and conserved residues in proteins. In this study, we included ESM-1v ^1^, ESM-2^2^, and ESM-C ^3^ for comparison, representing three successive generations of the ESM family of pLMs. The evolutionary plausibility of each variant is computed by following Eq. 3. To compute evaluation metrics including AUROC, AUPRC, and nDCG, we negated the ESM-predicted values so that higher scores correspond to greater evolutionary implausibility. To compute the F1 score, variants with *e*(***x***^MT^) *<* 0 were classified as SBI variants. For identifying functionally important residues, we computed the average ESM-predicted plausibility score across all possible single-point mutations at each residue and then applied a negation. Consequently, residues with higher scores correspond to sites whose mutations are generally less evolutionarily plausible, indicating stronger evolutionary constraints and suggesting greater functional importance. ESM-1v is an ensemble of five pre-trained models (esm1v_t33_650M_UR90S_{1, …, 5}). For ESM-2 and ESM-C, we chose their best-performing versions in terms of functional site identification on Human Domainome 1 (esm2_t30_-150M_UR50D and esmc_300m) as the representatives.

#### Stability-based predictors

We included FoldX ^4^ ΔΔ*G*, SPURS ^5^ ΔΔ*G*, and ESM-IF1^6^ as stability-based predictors for SBI variants and functional sites. Both FoldX ΔΔ*G* and SPURS ΔΔ*G* measure the change in the change in Gibbs free energy between the wild-type (WT) structure and the variant (MT) structure. When ΔΔ*G <* 0 kcal/mol, the variant is predicted to be stabilizing. For the computation of AUROC, AUPRC, and nDCG, we negate the value of ΔΔ*G* so that higher scores correspond to higher stability. ESM-IF1 is an inverse folding model designed to predict protein sequences based on their structures. It is pretrained to predict the amino acid at each residue in an autoregressive manner, conditioned on the surrounding sequence context. Unlike other sequence-based ESM models, however, ESM-IF1 incorporates the wild-type protein structure as an additional conditioning input. Studies have shown that the mutational effects predicted by ESM-IF1 correlate well with the measures of protein thermostability ^7;8^. Following Cagiada et al. ^9^, we compute the logodds ratio between the variant sequence and the wild-type sequence for mutational effect prediction (similar to Eq. 3), and a threshold of –7 is used, with variants scoring above this value considered to be stable. For functional site identification, guided by the concept of local energetic frustration ^10^, we compute the averaged predicted score across all possible single-mutations at each residue for all stability-based predictors and apply a negation for ΔΔ*G*. A higher residue score indicates that mutations at that residue tend to be stabilizing, suggesting energetic frustration around the residue and potential functional importance. For ESM-IF1, esm_-if1_gvp4_t16_142M_UR50 is used.

#### Functional model

Functional model ^11^ is a supervised ML model that combines sequence conservation models and biophysical models for classifying SBI variants and predicting functional sites. For each single-point mutation, the input features include an evolutionary conservation score predicted by GEMME ^12^, a ΔΔ*G* predicted by Rosetta ^13^, a hydrophobicity score, a contact number, and a chain neighbors averaged score. Each single-point mutation is classified into four different categories: WT-like, total-loss, SBI, and low abundance but high activity. To compute the AUROC of the Functional model for identifying SBI variants, we used the predicted probabilities of variants being classified as SBI as the input scores for evaluation. We implemented the Functional model using the Jupyter notebook provided by Cagiada et al. ^11^ on GitHub (https://github.com/KULL-Centre/_2022_functional-sites-cagiada).

#### Func-ESM

Func-ESM ^9^ is an unsupervised approach to disentangle mutation effects on protein function and stability using a constant axis-parallel classification boundary. Func-ESM leverages ESM-1b for variant effect prediction and ESM-IF1 for predicting variant effects on protein stability. Single-point variants that have an ESM-1b score smaller than -6.5 but an ESM-IF1 score larger than -7 are predicted to be SBI variants. Residues enriched with SBI variants are predicted to be functionally important residues. We implemented Func-ESM using the Jupyter notebook provided by Cagiada et al. ^9^ on GitHub (https://github.com/KULL-Centre/_2024_cagiada-jonsson-func).

#### Logistic regression model

Logistic regression model ^14^ uses sigmoidal curves to model the relationship between ESM-1v predicted fitness and stability measurements. The residuals from the fitted sigmoidal curve indicate whether mutations have larger or smaller effects on protein evolutionary plausibility than can be explained by changes in stability. Following Li et al. ^5^, we modeled the relationship between ΔΔ*G* predicted by FoldX ^4^ or SPURS ^5^ and ESM-1v predicted fitness using a sigmoidal curve. Similar to DETANGO, mutations with lower residues are predicted to be SBI variants. Score at each residue is the negation of the average scores of all single-mutations of this residue. Residues with higher function scores are predicted to be more functionally important.

#### Weighted average between ESM-1v and FoldX

We incorporated a weighted average between ESM-predicted evolutionary plausibility scores and FoldX-predicted stability changes (ΔΔ*G*) as a baseline for identifying functionally important residues annotated in the CDD. For each variant, the combined score is defined as *e*(***x***^MT^) + *η*ΔΔ*G*(***x***^MT^), where *η* is a tunable coefficient balancing the contributions of evolutionary and energetic terms. We varied the parameter *η* from 0 to 1 in increments of 0.1. For each residue, its predicted score is computed as the negative mean of the scores across all its single-point mutations. A higher residue score indicates that its mutations are evolutionarily unfavorable yet enriched with stabilizing variants, suggesting local energetic frustration and potential functional importance.

#### Linear regression and MLP regression

In addition to the Func-ESM ^9^ and the Logistic regression model ^5;14^, we introduced two baseline approaches that disentangle mutation effects on protein function and stability using classification boundaries. Linear regression models the relationship between ESM-1v and ΔΔ*G* using a linear boundary. MLP regression models the relationship between ESM-1v and ΔΔ*G* using a non-linear decision boundary learned from data. Residuals are defined as the proportion of the ESM-1v fitness not explained by the stability components. The lower the residuals for mutations, the more likely they are SBI variants. Score at each residue is the negation of the average scores of all single-mutations of this residue. Residues with higher function scores are predicted to be more functionally important. For a fair comparison with DETANGO, MLP regression was an ensemble of 10 replicas under a 10-fold random split. Both linear regression and MLP regression were implemented using the default parameters in the Python package Sklearn.

#### SeqIC

SeqICs are calculated for proteins from homologous families using information theory concepts ^15^. For each aligned position in the MSA, SeqIC is defined as the maximum possible entropy minus the observed entropy, based on the distribution of amino acids. A higher SeqIC indicates higher sequence conservation at the aligned position, thereby more likely to be critical to the protein’s function.

### A.2 Additional implementation details of DETANGO

#### Hyperparameter selection

DETANGO is an unsupervised framework designed to disentangle the effects of mutations on protein stability and function as predicted by pLMs. Since it is not trained against explicit functional labels, all parameters and hyperparameters in the DETANGO framework were optimized to minimize an unsupervised, function-irrelevant disentanglement loss:

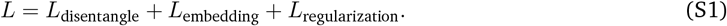

The hyperparameter search space, including learning rate, batch size, number and dimension of hidden layers in the prediction heads (i.e., *g*_structural_, *g*_functional_, and *g*_stability_), number and dimension of layers in the projection networks (i.e., *g*_projection_), and dropout rate, is summarized in Supplementary Table 4. Optimal hyperparameters were selected by minimizing the validation loss, averaged over ten independent replicas trained on four representative proteins from Cagiada et al. ^11^ (Supplementary Table 1; Supplementary Fig. 7).

#### Using abundance to guide disentanglement

Compared to Func-ESM ^9^ and Logistic regression model ^5;14^, a key advantage of DETANGO is its flexibility in leveraging different forms of stability measurements to guide the disentanglement of evolutionary plausibility, even when such measurements are available only for a subset of single-point mutations. By employing a lightweight MLP to model the relationship between stability measurements and the structural plausibilities of protein variants, DETANGO makes no prior assumptions about this relationship and instead learns it directly in a data-driven manner. When training with incomplete stability measurements, where only a subset of single-point mutations has available data, we applied a masking scheme to compute the stability-prediction loss only for those mutations with known measurements during embedding decomposition:

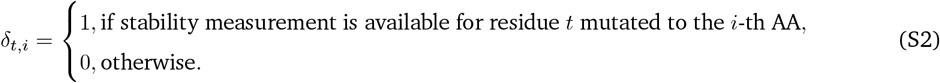

When computing the objective function following Eq. S1, only predictions with available ground-truth labels contribute to the loss terms *L*_disentangle_ and *L*_stability_. Specifically, under the masking scheme, the loss computation for each minibatch is formulated as:

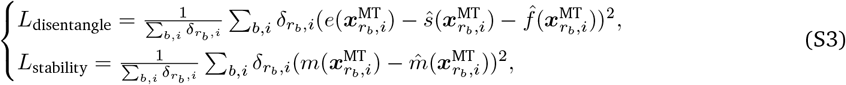

where *r*_*b*_ denotes the *b*-th protein residue in the batch, and 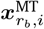 denotes the variant with a substitution of the *r*_*b*_-th residue to the *i*-th AA from the alphabet.

#### Mutual information-based loss function for embedding disentanglement

We further experimented with the incorporation of a mutual information-based loss into the training objective of DETANGO to promote the embedding disentanglement and reduce the dependence between the function-specific representation **h**^(*f*)^ and the stability-specific representation **h**^(s)^. We incorporated a loss function *L*_MI_ to minimize the mutual information between **h**^(*f*)^ and **h**^(s)^

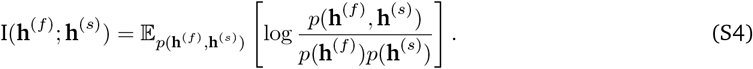

We adopt the variational Contrastive Log-ratio Upper Bound (vCLUB) framework ^16^ as an estimation of the mutual information between **h**^(*f*)^ and **h**^(s)^Assuming the joint distribution of **h**^(*f*)^ and **h**^(s)^is intractable and the conditional relationship between the two variables (e.g., *p* **h**^(s)^**h**^(*f*)^)) is unknown, vCLUB uses a variational distribution *q*_*θ*_ (**h**^(s)^ | **h**^(*f*)^) to approximate the conditional distribution and is defined as

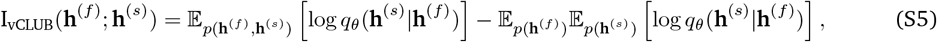

where *q*_*θ*_(**h**^(s)^|**h**^(*f*)^) is approximated with a multilayer perceptron parameterized with *θ*. With sample pairs 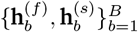 in a batch, I_vCLUB_(**h**^(*f*)^; **h**^(s)^) has an unbiased estimation as:

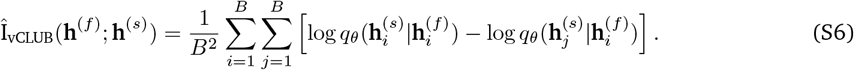

We introduced an additional loss function term *L*_mi_ that equals to Î_vCLUB_(**h**^(*f*)^; **h**^(s)^) as a regularizer in DETANGO. The overall loss function for training DETANGO (MI) is a weighted summation of the four loss functions:

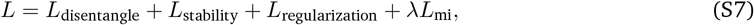

where *λ* is set to 0.1 to ensure that the value of *L*_mi_ is on the same scale as the other three loss function terms.

When benchmarking DETANGO (MI) against baseline approaches and the standard DETANGO for identifying CDD-annotated functional sites in Human Domainome 1^14^ proteins, the inclusion of the MI–based loss term yielded modest but statistically non-significant performance gains (Supplementary Fig. 6). Accordingly, this loss term was not incorporated into the final DETANGO framework.

## B Supplementary Figures

**Supplementary Figure 1.**
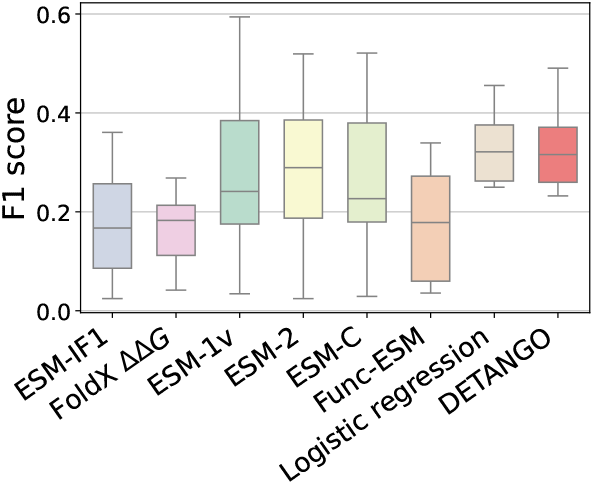
DETANGO achieves accurate classification for stable-but-inactive (SBI) variants. DETANGO was compared with seven unsupervised baseline approaches (ESM-IF1, FoldX ΔΔ*G*, ESM-1v, ESM-2, ESM-C, Func-ESM, Logistic regression model) for SBI classification on 11 proteins, using F1 score as the evaluation metric. The middle line of the box plot indicates the median of the data, while the box extends from the first quartile to the third quartile. The whiskers reach the smallest and largest values that fall within 1.5 times the interquartile range beyond the first and third quartiles.

**Supplementary Figure 2.**
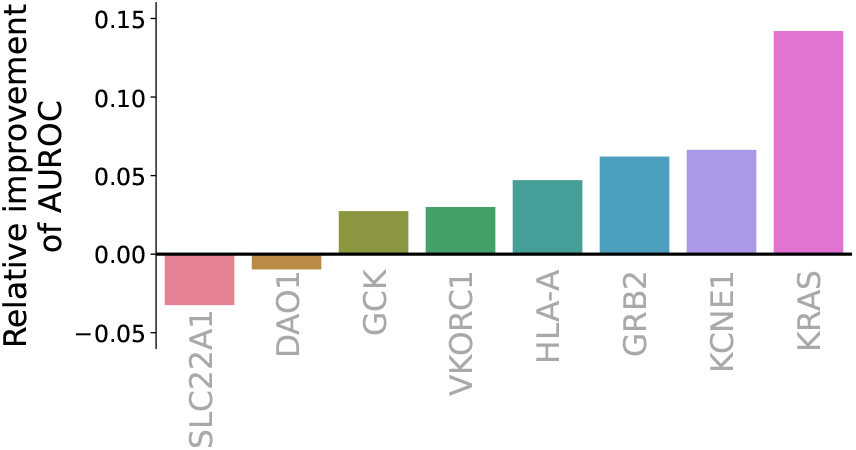
DETANGO surpasses Functional Model for classifying SBI variants. DETANGO was evaluated against a supervised ML baseline (Functional Model) for SBI classification across eight proteins that were not included in the training of Function Model, using the relative improvement of DETANGO over Function Model in AUROC as the evaluation metric. The Functional Model had been trained exclusively on variants from NUDT15, PTEN, and CYP2C9.

**Supplementary Figure 3.**
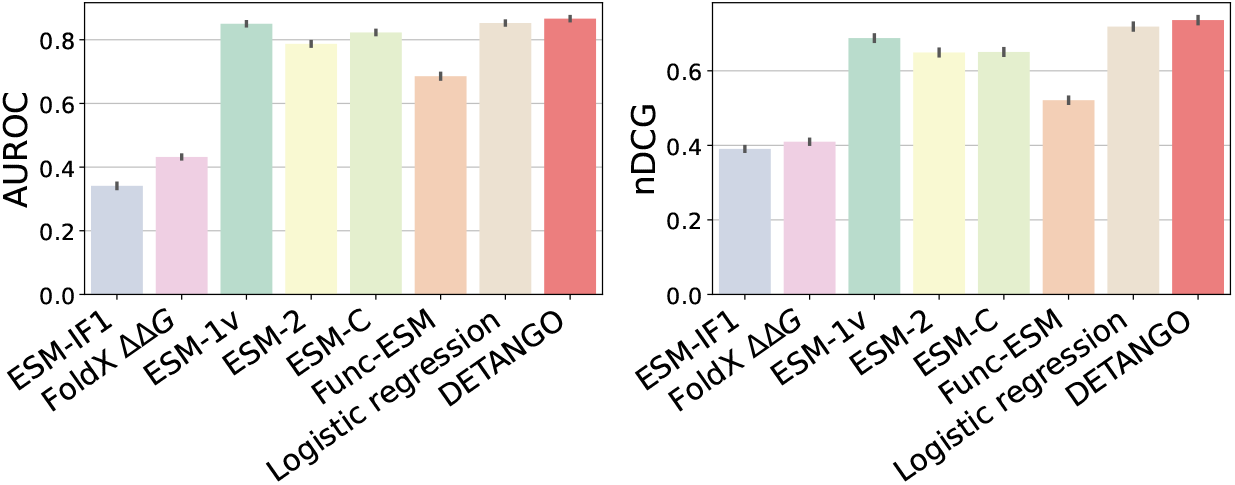
DETANGO accurately identifies functionally important residues in Human Domainome 1 proteins. DETANGO was compared with seven unsupervised baseline approaches (ESM-IF1, FoldX ΔΔ*G*, ESM-1v, ESM-2, ESM-C, Func-ESM, and Logistic regression model) for identifying CDD-annotated functional residues in proteins from Human Domainome 1, using AUROC and nDCG as the evaluation metrics.

**Supplementary Figure 4.**
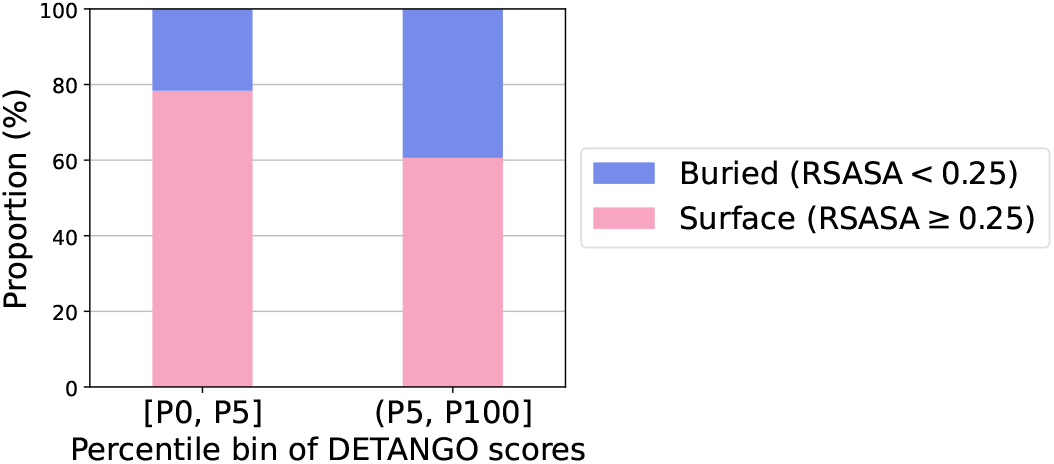
Solvent accessibility of residues in Human Domainome 1 proteins ranked by DETANGO. The proportions of surface residues were compared between residues within the bottom DETANGO score percentile ([P0, P5]) and all other residues ((P5, P100]). Surface residues were defined as those with a relative solvent-accessible surface area (RSASA) of no less than 0.25.

**Supplementary Figure 5.**
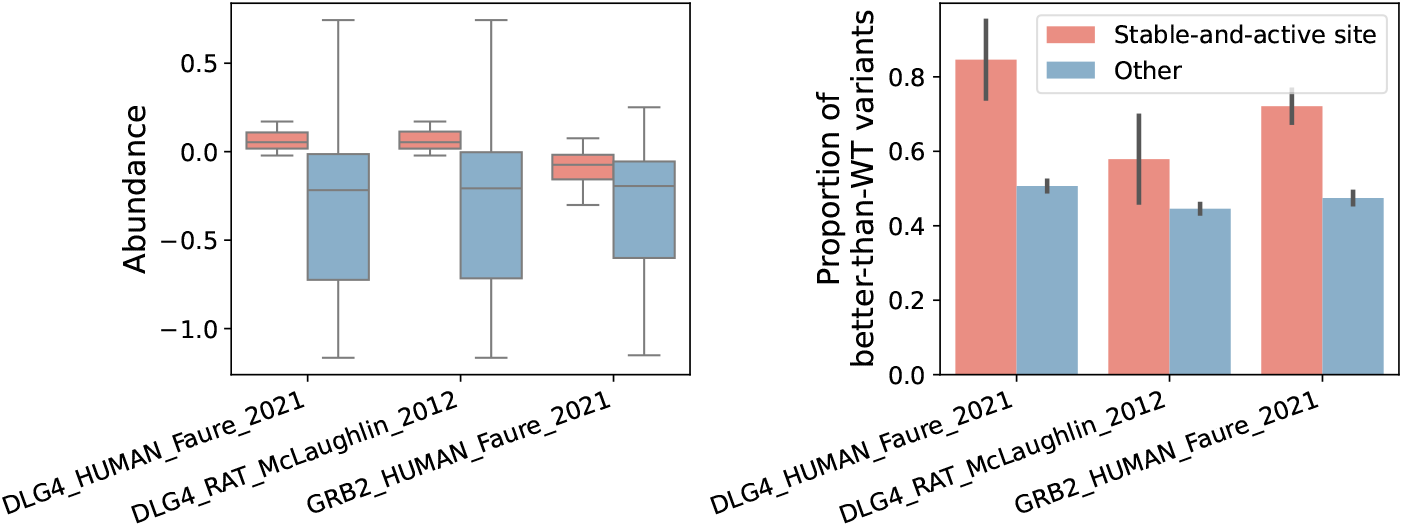
Comparison of abundance and functional readouts at stable-and-active sites. The abundances and functional readouts of single-point mutations were compared between stable-and-active sites and other residues. Abundance data were retrieved from Human Domainome 1.0, and functional data were collected from ProteinGym v1.0. Stable-and-active sites were defined as residues with DETANGO scores in the lowest 10% for each protein. In the box plot, the central line represents the median, boxes indicate the interquartile range (IQR), and whiskers extend to the most extreme data points with 1.5*×*IQR. Error bars in the bar plot indicate the standard error (SE) of the mean

**Supplementary Figure 6.**
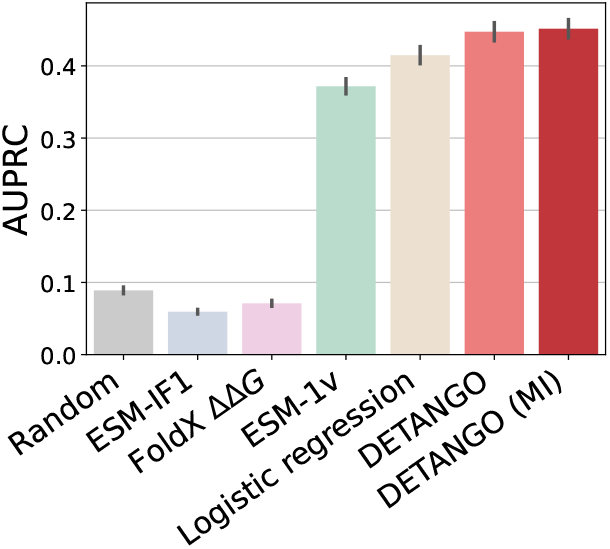
Impact of Mutual Information (MI)–Based Regularization on Functional Residue Identification. DETANGO (MI) is a variant of DETANGO that introduces an MI-based regularization loss to enhance the disentanglement of structural and functional embeddings within the DETANGO framework. DETANGO (MI) was compared with DETANGO and five unsupervised baseline approaches (random, ESM-IF1, FoldX ΔΔ*G*, ESM-1v, Func-ESM, and logistic model) for identifying CDD-annotated functional residues in proteins from Human Domainome 1, using AUPRC as the evaluation metric.

**Supplementary Figure 7.**
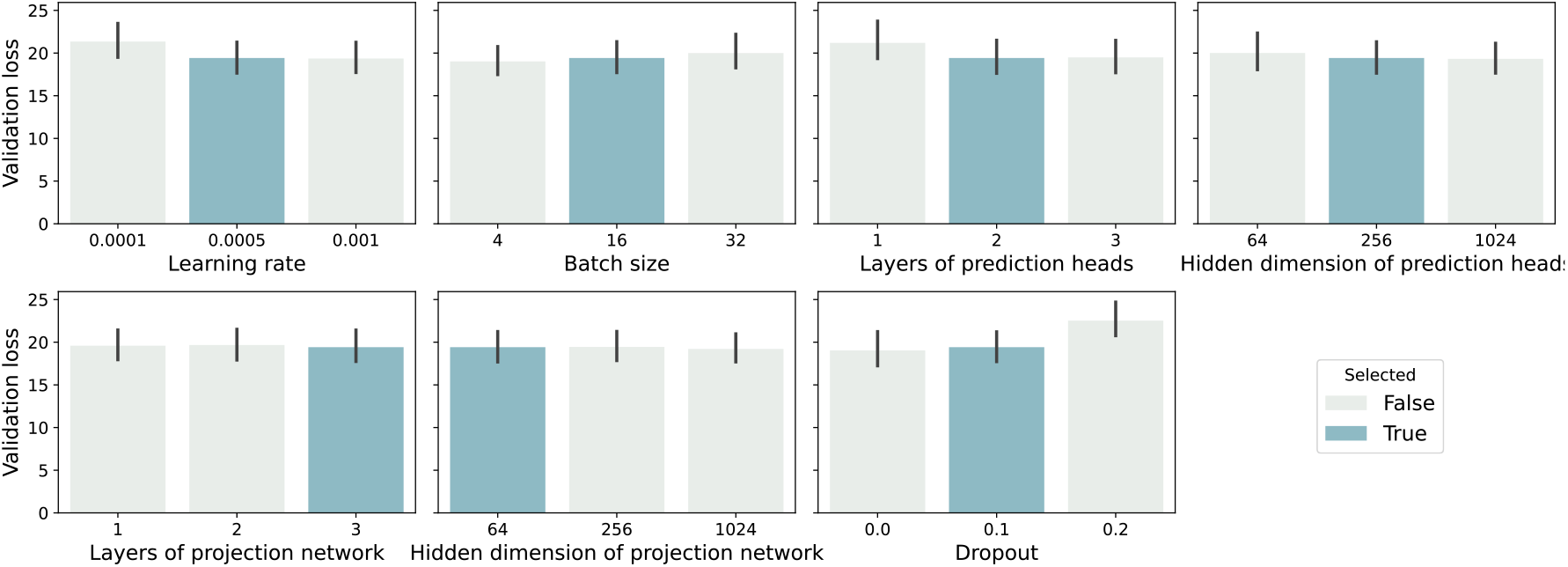
Robustness of Hyperparameter Selection in the DETANGO Framework. The hyperparameters of DETANGO were optimized to minimize the validation loss averaged over ten independent replicas on four representative proteins from Cagiada et al. ^11^, including CYP2C9, NUDT15, PTEN, and GRB2. The results demonstrate that the model’s performance remains robust across a broad range of hyperparameter settings.

## C Supplementary Tables

**Supplementary Table 1.** Description of the dataset used for stable-but-inactive (SBI) variant classification. Multidimensional deep mutational scanning (DMS) assays for 11 proteins were collected from Cagiada et al. ^11^ and ProteinGym v1.0^7^ for benchmarking.

| Gene Name | UniProt ID | Assay types | Data source |
| --- | --- | --- | --- |
| CYP2C9 | P11712 (CP2C9_HUMAN) | Abundance, Enzyme activity | Cagiada et al.<sup>11</sup> |
| NUDT15 | Q9NV35 (NUD15_HUMAN) | Abundance, Cytotoxicity | Cagiada et al.<sup>11</sup> |
| PTEN | P60484 (PTEN_HUMAN) | Abundance, Lipid phosphatase activity | Cagiada et al.<sup>11</sup> |
| GRB2 | P62993 (GRB2_HUMAN) | Abundance, Binding | Cagiada et al.<sup>11</sup> |
| GCK | P35557 (HXK4_HUMAN) | Abundance, Enzyme activity | ProteinGym v1.0<sup>7</sup> |
| KCNE1 | P15382 (KCNE1_HUMAN) | Expression, Channel function | ProteinGym v1.0<sup>7</sup> |
| DAO1 | P80324 (OXDA_RHOTO) | Expression, Enzyme activity | ProteinGym v1.0<sup>7</sup> |
| HLA-A | Q53Z42 (Q53Z42_HUMAN) | Expression, Binding | ProteinGym v1.0<sup>7</sup> |
| KRAS | P01116 (RASK_HUMAN) | Abundance, Binding | ProteinGym v1.0<sup>7</sup> |
| SLC22A1 | O15245 (S22A1_HUMAN) | Abundance, Cytotoxicity | ProteinGym v1.0<sup>7</sup> |
| VKORC1 | Q9BQB6 (VKOR1_HUMAN) | Abundance, Activity | ProteinGym v1.0<sup>7</sup> |

**Supplementary Table 2.** Description of the mammalian *α* and *β* globin subfamilies. Following Freiberger et al. ^15^, the functional patterns of 19 *α* globins and 20 *β* globins were analyzed. ^*†*^ P01972 was excluded due to a sequence mismatch between its UniProt entry and the corresponding AlphaFold DB (v4) model.

| Organism | $\alpha$ globin | $\beta$ globin |
| --- | --- | --- |
| *Chrysocyon brachyurus* (Maned wolf) | P60523 | P60526 |
| *Bos taurus* (Bovine) | P01966 | P02070 |
| *Odocoileus virginianus virginianus* (Virginia white-tailed deer) | P01972<sup>†</sup> | P02074 |
| *Sus scrofa* (Pig) | P01965 | P02067 |
| *Equus asinus* (Donkey) | P01959 | D1MPT0 |
| *Equus caballus* (Horse) | P01958 | P02062 |
| *Homo sapiens* (Human) | P69905 | P68871 |
| *Ovis aries* (Sheep) | P68240 | P02075 |
| *Oryctolagus cuniculus* (Rabbit) | P01948 | P02057 |
| *Cavia porcellus* (Guinea pig) | P01947 | P02095 |
| *Bubalus bubalis* (Domestic water buffalo) | Q9TSN8 | P67820 |
| *Capra hircus* (Goat) | P0CH25 | P02077 |
| *Felis catus* (Cat) | P07405 | P07412 |
| *Rattus norvegicus* (Rat) | P01946 | P02091 |
| *Pteropus vampyrus* (Large flying fox) | D0VX09 | D0VX08 |
| *Camelus dromedarius* (Arabian camel) | P63106 | P68231 |
| *Canis lupus familiaris* (Dog) | P60529 | P60524 |
| *Lepus europaeus* (European hare) | F2Z285 | P08535 |
| *Mammuthus primigenius* (Siberian woolly mammoth) | D3U1H8 | D3U1H9 |
| *Helogale parvula* (Common dwarf mongoose) | A0A0R4I977 | A0A0R4I978 |

**Supplementary Table 3.** Description of the human RAS superfamily. Following Freiberger et al. ^15^, the functional patterns of 148 human proteins from the RAS superfamily were analyzed: RAS (n=37), RAB (n=63), RHO (n=22), and ARF (n=26). ^*†*^ Q9BR65 and Q6PJZ0 were excluded from the analyses because their corresponding UniProt entries have been merged into P01112 and Q9H0T7, respectively.

| Family | UniProt ID | Family | UniProt ID | Family | UniProt ID | Family | UniProt ID |
| --- | --- | --- | --- | --- | --- | --- | --- |
| RAS | Q9H628 | RAB | Q9NRW1 | RAB | Q9UL26 | RHO | Q9H4E5-1 |
| RAS | Q99578-1 | RAB | Q9H0N0 | RAB | Q9H0T7 | RHO | P17081 |
| RAS | P61225 | RAB | P51149 | RAB | Q9NP72 | RHO | P61587 |
| RAS | P62834 | RAB | Q96AH8 | RAB | A4D1S5-1 | RHO | P52198 |
| RAS | P01111 | RAB | O14966 | RAB | C9JJQ5 | RHO | O00212 |
| RAS | Q8IYK8 | RAB | Q9NP90 | RAB | P62820-1 | RHO | Q9HBH0-1 |
| RAS | P11233 | RAB | Q92930 | RAB | Q9H0U4 | RHO | O94844 |
| RAS | Q96A58 | RAB | Q9BW83-2 | RAB | Q6FIG4 | RHO | Q92730 |
| RAS | P55042 | RAB | Q8IZ41-1 | RAB | Q6PJZ0<sup>†</sup> | RHO | Q96L33 |
| RAS | Q96S79 | RAB | P20340-1 | RAB | A8MSP2 | RHO | Q7L0Q8-1 |
| RAS | Q8TAI7-1 | RAB | P61006 | RAB | Q7Z4W7 | ARF | P40617 |
| RAS | O95057 | RAB | P20340-2 | RAB | Q14964 | ARF | Q6T311 |
| RAS | Q9BR65<sup>†</sup> | RAB | P61026 | RAB | P61020 | ARF | P40616 |
| RAS | Q7Z444 | RAB | P51148 | RAB | P20339 | ARF | P36406-1 |
| RAS | P55040 | RAB | Q969Q5 | RAB | P61018-1 | ARF | P84077 |
| RAS | Q6T310 | RAB | P57735 | RAB | Q53GC2 | ARF | P61204 |
| RAS | Q9BPW5 | RAB | Q9ULW5 | RAB | Q86YS6 | ARF | P18085 |
| RAS | P01116-1 | RAB | P51159-1 | RAB | Q96S21 | ARF | P84085 |
| RAS | O95661 | RAB | O00194 | RAB | Q12829 | ARF | P62330 |
| RAS | P11234 | RAB | P51157-2 | RAB | Q8WXH6 | ARF | P36404 |
| RAS | P01112 | RAB | Q9ULC3 | RAB | O95716 | ARF | Q13795 |
| RAS | Q9Y3L5 | RAB | P61019 | RAB | Q96E17 | ARF | P36405 |
| RAS | P10114 | RAB | Q15771 | RAB | P20337 | ARF | Q9Y6B6 |
| RAS | Q9NYS0 | RAB | Q13637 | RAB | P20336 | ARF | Q9NVJ2 |
| RAS | P01116-2 | RAB | Q14088 | RAB | P51151 | ARF | Q9NXU5 |
| RAS | Q96HU8 | RAB | Q9H082 | RAB | Q96DA2 | ARF | Q9NR31 |
| RAS | Q9NYN1 | RAB | Q9BZG1-1 | RHO | Q15669 | ARF | Q8N4G2 |
| RAS | Q9Y272 | RAB | Q15286 | RHO | P60953-1 | ARF | P36406-2 |
| RAS | P61224 | RAB | Q8WUD1 | RHO | P60953-2 | ARF | P36406-3 |
| RAS | Q96D21 | RAB | O95755-1 | RHO | P63000-2 | ARF | Q969Q4 |
| RAS | Q9NYR9-1 | RAB | Q53EY4 | RHO | P63000-1 | ARF | Q96BM9 |
| RAS | B7Z7H6 | RAB | Q9UL25 | RHO | P15153 | ARF | Q9H0F7 |
| RAS | O14807 | RAB | P62491 | RHO | P60763 | ARF | Q9Y689 |
| RAS | Q92963 | RAB | Q15907 | RHO | P61586 | ARF | Q96KC2 |
| RAS | Q92737-1 | RAB | Q6IQ22 | RHO | P84095 | ARF | Q3SXY8-1 |
| RAS | P10301 | RAB | P51153 | RHO | P62745 | ARF | Q0P5N6 |
| RAS | Q15382 | RAB | P61106 | RHO | Q9BYZ6 |  |  |
| RAS | O75628 | RAB | P59190-2 | RHO | P08134 |  |  |

**Supplementary Table 4.** Search space of hyperparameters used in the DETANGO framework. The table summarizes the ranges of hyperparameters explored during model optimization, including learning rate, batch size, number and dimension of hidden layers in the prediction heads and projection network, and dropout rate. These hyperparameters were tuned to minimize the unsupervised disentanglement loss on the validation set.

| Parameter | Search space | Selected value |
| --- | --- | --- |
| Learning rate | [1e-4, 5e-4, 1e-3] | 5e-4 |
| Batch size | [4, 16, 32] | 16 |
| Number of layers in the prediction heads | [1, 2, 3] | 2 |
| Dimension of layers in the prediction heads | [64, 256, 1024] | 256 |
| Number of layers in the projection network | [1, 2, 3] | 3 |
| Dimension of layers in the projection network | [64, 256, 1024] | 64 |
| Dropout | [0.0, 0.1, 0.2] | 0.1 |

